# Novel cell- and stage-specific transcriptional signatures defining *Drosophila* neurons, glia and hemocytes

**DOI:** 10.1101/2022.06.30.498263

**Authors:** Rosy Sakr, Pierre B. Cattenoz, Alexia Pavlidaki, Laura Ciapponi, Marta Marzullo, Nivedita Hariharan, Tina Mukherjee, Angela Giangrande

## Abstract

Cell types can be now defined at unprecedented resolution using high throughput assays. We analyzed the transcriptional signatures of *Drosophila* neurons, glia and hemocytes, as examples of cell types that are related by position (glia/neurons) or function (glia/hemocytes) or that are unrelated (neurons/hemocytes). The most related cells display the highest similarity level (neurons and glia), the least related ones, the lowest (neurons and hemocytes), however, cells can show plastic features. Glia are much more similar to neurons than to hemocytes in the embryo, but are equally similar to the two cell types in the larva, when hemocytes acquire more immune functions. Larval glia and hemocytes display common as well as specific immune features, such as the glia-specific NimA receptor, in agreement with the different environment faced by each cell types. Surprisingly, time represents a key identity parameter, as neurons, hemocytes and glia group more significantly by the stage than by the cell type and larval cells show upregulation of genes involved in chromatin organization and in DNA repair. This latter group of genes is linked to changes in gene expression levels and chromatin organization, revealing a function of these genes beyond DNA repair. Finally, the metabolic profiles reveal cell type-specific signatures and an overall shift from an embryonic, anabolic state to a larval, catabolic state.

## Introduction

Traditionally, cell types have been defined by morphology, position, origin and function, relying on the expression of few proteins. While the combination of these parameters has produced valuable information, each of them on their own can lead to misleading conclusions. Cells in different organs can display similar functions and cells within the same organ can have different properties. Moreover, the *a priori* selection of proteins can introduce bias and neglect the wealth of molecular information intrinsic to each cell type. Finally, functional plasticity and dynamic states may conduct to a reductionist classification. To ensure a more unbiased and quantitative analysis, we used a high throughput approach and a data-driven computational method on neurons, hemocytes and glia. To account for plasticity, we analyzed them at two stages, embryo and larva.

*Drosophila* neurons and glia differentiate from common, ectodermal, precursors. Neurons sense, process and transmit information or secrete neuropeptides (Garces and Thor, 2006; Landgraf and Thor, 2006; Schmid et al., 1999). Glia are involved in nervous system development, function, and maintenance. They control cell proliferation (Ebens et al., 1993), axonal and synapse development (Brink et al., 2012; Ou et al., 2014), neuronal insulation and survival (Shepherd, 2000; Stork et al., 2008; Volkenhoff et al., 2015). They also establish the blood-brain barrier (BBB) (Abbott, 2005) and provide the immune function within the nervous system (Awasaki and Ito, 2004; Sonnenfeld and Jacobs, 1995a; Watts et al., 2004). The mesodermally derived hemocytes patrol the organism to ensure cellular immunity outside the nervous system (Lebestky et al., 2000; Lemaitre et al., 1996; Tepass et al., 1994), hence sharing the function but not the origin with glial cells, while sharing neither the origin nor the function with neurons.

Transcriptome comparison highlights cell-specific features. Neurons and hemocytes show the least related transcriptional landscapes regardless of the stage. Glia and neurons stably share the highest similarity, in line with the fact that they constantly interact to ensure the homeostasis of the nervous system. Glia and hemocytes display significant similarity in the larva. This goes along with a significant change of the hemocyte transcriptional profile from the embryo to the larva, an open system that has to react to external stimuli as pathogens. Although larval glia and hemocytes share an immune function, the two cell types encounter distinct environments, pathogens for hemocytes, axon and dendrite debris for glia. Accordingly, the Eater and the Nimrod C1 (NimC1) scavenger receptors are specific to hemocytes, NimA is specific to glia.

Similarly, some metabolic features are common, others are specific with respect to tissue and stage. The key glycolytic gene *lactate dehydrogenase (ldh)* is upregulated in all the three cell types in the embryo, down-regulated in the larva. The expression of lipogenic genes, on the other hand, is upregulated in all three embryonic cell types, but changes discretely as the embryo transitions to the larval stage. Hemocytes dramatically shift to a lipolytic state while neurons maintain their lipogenic state. Larval glia maintain the expression of lipogenic genes but shows additional upregulation of lipolytic pathway genes. The observed common metabolic states may be a reflection of the overall developmental program, but the cell-specific states may highlight metabolic needs necessary to accommodate distinct functional roles.

Transcriptome comparison also highlights common stage-specific features. Larval cells express genes controlling chromatin organization as well as DNA repair at high levels. One of the main DNA repair genes is *rad50,* involved in DNA double strand break (DSB) repair. Its depletion triggers a genome-wide change in chromatin organization and mis-regulation of gene expression, pointing to a role beyond DSB. Finally, the three cell types display a glycolytic and lipogenic state in the embryo, a lipolytic and oxidative state in the larva, changes that are likely linked to external inputs such as feeding and respiration.

Altogether, our genome-wide analysis makes it possible to disentangle the relative importance of origin, position, function and time in the definition of cell types. The expression profile of a cell population is strongly affected by their origin in the embryo, while larval cells acquire additional levels of specialization and become more affected by the function. Moreover, the three cell types display different degrees of plasticity, the migratory hemocytes exposed to the external challenges being the most plastic ones. This work paves the way for systematic identification of gene networks and expression programs, which will ultimately shed light on the biological mechanisms characterizing cell diversity in multicellular organisms.

## Results

### Impact of origin, stage and function on the transcriptional landscapes

To assess the impact of origin, function and position on the definition of a cell type, we compared the transcriptomes of neurons and glia, which share the origin; those of hemocytes and glia, which share the scavenging activity; and those of neurons and hemocytes, which share neither origin nor function. We performed the comparison in mature embryos (stage 16 or E16) and in wandering third instar larvae (L3), to understand whether expression profiles change from a small, closed and immobile system to a bigger, open system that is able to grow and to respond and adapt to the environment. Glia, hemocytes and neurons were purified from transgenic lines specific to each cell type: *repo-nRFP, srp(hemo)Gal4/+; UAS-RFP/+* and *elav-nRFP*, respectively. The hierarchical clustering of the transcriptomes shows a good correlation between the biological replicates, highlighting a low biological variability (**Figure 1-a**).

**Figure 1:**
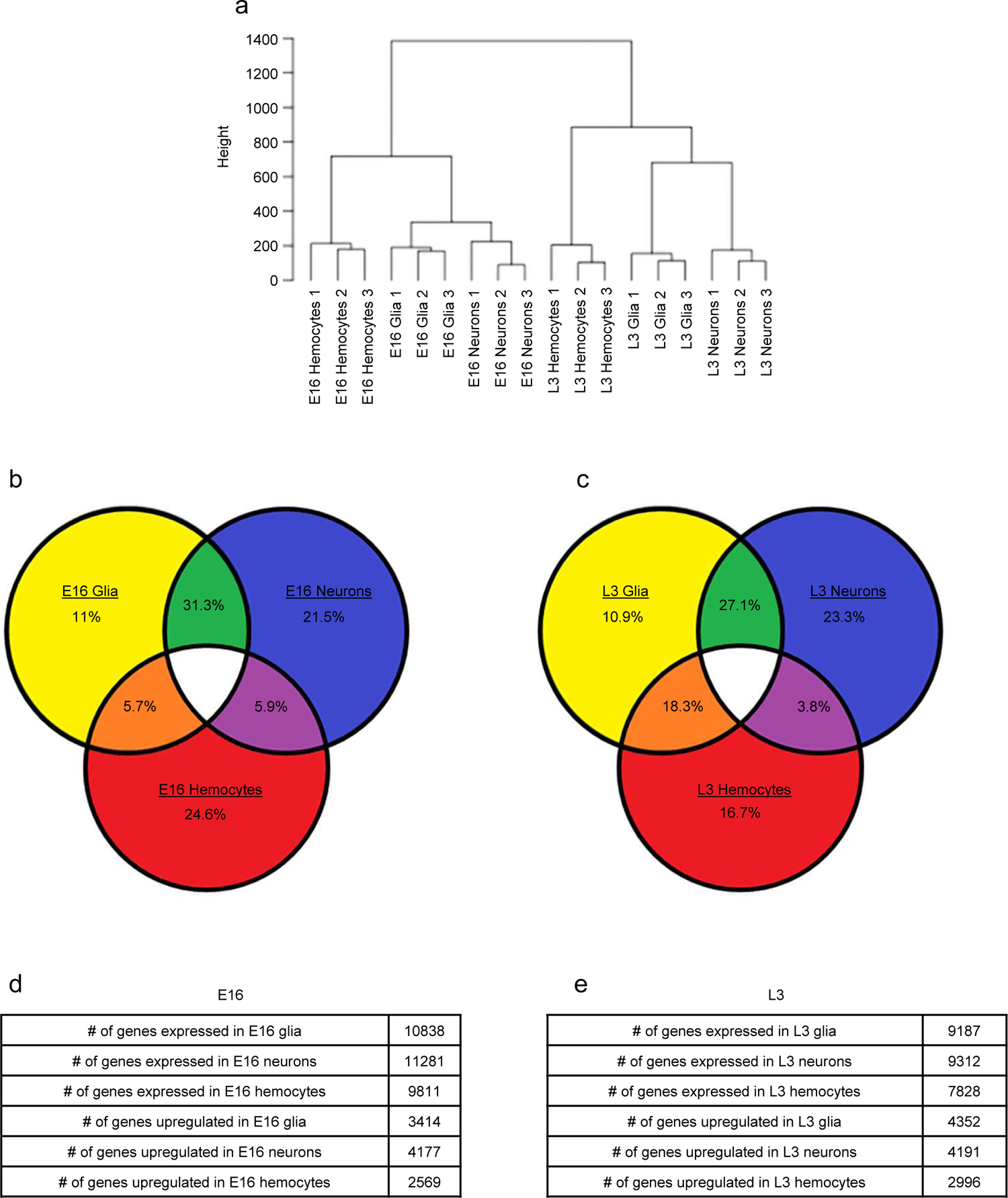
RNAseq analysis in embryonic and larval glia, neurons and hemocytes. (a) Dendrogram showing the hierarchical clustering based on the distance between samples. The height indicates the distance. Euclidean distance, Ward criterion were used for this representation. (b-c) Venn diagrams showing the distribution of differentially expressed genes between neurons (blue), glia (yellow) and hemocytes (red) at E16 (b) and L3 (c). Since only upregulated genes are used for the analysis, the intersection between all three cell types is empty (white). The percentage of genes in each category is indicated in the diagram. The percentage was calculated based on all genes used for the analysis. (d,e) The total number of genes expressed (top three lines) and upregulated compared the other two cell types (bottom three lines) at E16 (d) and L3 (e).

The developmental stage is the main factor of heterogeneity, more important than the cell type, as the highest correlation in the dendrogram is seen for samples of the same stage. Thus, each cell type is more similar to the two others at the same stage than to the same cell type at a different stage. The dendrogram also shows that glia and neurons group together, sharing the highest degree of similarity. We next performed pairwise comparisons of the number of differentially expressed genes to evaluate the commonalities and the differences between neurons, glia and hemocytes. We defined significantly upregulated genes based on transcripts that are enriched in one population compared to the other following three parameters: number of reads > 15, adjusted p-value <0.05 and fold change >1.5 between the number of reads. Upregulated genes in each cell type were compared in the Venn diagram. The comparison of the cell-specific transcriptomes is shown in (**Figure 1-b**) for the embryo, in (**Figure 1-c**) for the larva. The highest number of shared upregulated genes concerns E16 neurons and glia (in green) (31.3% of all genes included in the diagram), while only 5.7% of shared upregulated genes are found between E16 hemocytes and glia (in orange). Interestingly, the shared enriched transcripts between hemocytes and glia reach 18.3% by the larval stage, suggesting that glia and hemocytes acquire common molecular features by L3, despite their distinct origin. Finally, the Venn diagram and the dendrogram indicate that glia and neurons maintain a high degree of similarity in the larva (27.1%). Neurons and hemocytes remain the most unrelated cell populations, with 5.9% of shared enriched transcripts at E16, 3,8% at L3.

Interestingly, the number of expressed genes is overall lower in L3 compared to E16, and this is true for all cell types (transcripts with at least 15 reads, **Figure 1-d,e, top three lines**). By contrast, the number of upregulated genes in each cell type increases by L3 (**Figure 1-d,e, bottom three lines).** 10838 genes are expressed in E16 glia, 11281 in E16 neurons and 9811 in E16 hemocytes. By L3, 9187 genes are expressed in glia, 9312 in neurons and 7828 in hemocytes, which means that at least 1500 genes are only expressed at E16 in each cell type compared to L3 (**Figure 1-d,e)**. This is not due to a low sequencing depth, since the same number of cells was used for each sample and there are no differences between the number of reads mapped for embryonic and larval cells (**Sup. Figure 1-a**). To further understand the molecular mechanisms underlying the stage differences, we analyzed the expression profiles of transcription factors, proteins that coordinately control downstream effector genes. In line with the increased number of cell-specific upregulated genes in the larva, the number of transcription factors commonly expressed in all three cell types decreases by L3 (**Sup. Table 1-a,b).** Thus, differentiated cells become more specialized over time.

In sum, our results show that the developmental stage has a stronger impact on the transcriptome than the cell type. In agreement with their respective origin and position, glia and neurons display the strongest similarity, hemocytes and neurons display the opposite behavior. Glia and hemocytes become more similar by L3. Finally, larval differentiated cells express fewer genes than the embryonic counterparts, but, proportionally, more genes that are upregulated in a cell-specific manner.

### Cell-specific signatures in the embryo define neurons and glia but not hemocytes

The comparison of the transcriptomes indicates that almost half of the transcripts are common to neurons, glia and hemocytes (see **Figure 1-d,e**: number of genes expressed – number of genes upregulated in each cell type), calling for more refined parameters to define the three cell types. Since a significant fraction is enriched only in one cell type or shared by two cell populations, we predicted that a pairwise comparison associated with quantitative and Gene Ontology (GO) analyses would allow the disclosure of the signatures and functions that better define cell types.

In the embryo, 1649 (out of 11281) genes are upregulated in neurons compared to glia (in blue), 992 in glia compared to neurons (out of 10838) (in yellow) (**Figure 2–a**). Thus, relatively few genes are upregulated in a specific cell type. The vast majority of the genes are non-differentially expressed, with number of reads > 15 and adjusted p-value of fold change > 0.05. On average, these genes are expressed at lower levels than those upregulated in a specific cell type (**Sup. Figure 1-b,c**). Amongst the non-differentially expressed genes, the ones with fold change < 1.5 were considered as commonly expressed, mostly involved in biological processes found in all cells such as ncRNA processing, ribosome biogenesis and mRNA metabolic process (**Sup. Figure 1-d, Sup. Table 2-a**). The transcripts enriched in neurons are mostly involved in neuronal pathways, such as synaptic transmission, neurotransmitter secretion and neuron projection guidance (**Figure 2-b, Sup. Table 2-b)**. Those enriched in glia are involved in the establishment of the BBB, axon ensheathment and the development of glial cells (**Figure 2-c, Sup. Table 2-c).**

**Figure 2:**
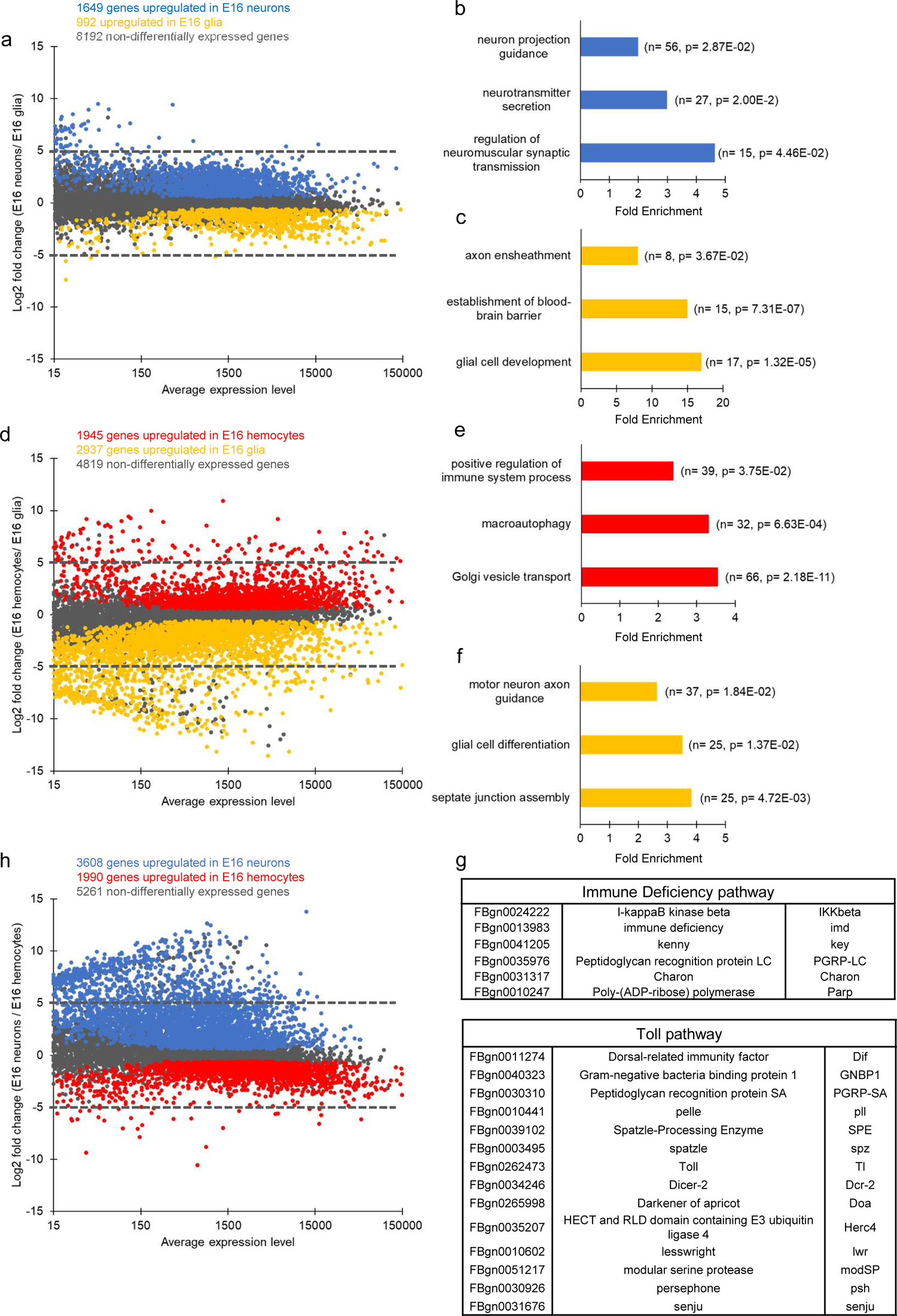
Pairwise transcriptome comparisons in the embryo. (a) Transcriptome comparison of E16 neurons and glia, MA plot. The x-axis reports the average gene expression levels, the y-axis, the log2 fold change between the genes in E16 neurons/E16 glia. Genes significantly upregulated in E16 neurons are shown in blue, genes significantly upregulated in E16 glia are shown in yellow and non-differentially expressed genes are shown in gray. (b,c) Gene Ontology (GO) term enrichment analysis in E16 neurons (blue) and E16 glia (yellow). The fold enrichment for a subset of representative GO terms is displayed on the x-axis, the number of genes within the GO term and the P-value of the GO term enrichment are indicated in brackets. The complete results of the GO term analysis are shown in **Sup. Table 2-b,c.** (d) Transcriptome comparison of E16 hemocytes and glia. MA plot as in (a), the y-axis reports the log2 fold change E16 hemocytes/E16 glia. Genes significantly upregulated in E16 hemocytes are shown in red, genes significantly upregulated in E16 glia are shown in yellow and non-differentially expressed genes are shown in gray. (e,f) GO term enrichment analysis in E16 hemocytes (red) and E16 glia (yellow). The fold enrichment for a subset of representative GO terms is displayed on the x-axis, the number of genes and the P-value of the GO term enrichment are indicated in brackets. The complete results of the GO term analysis are shown in **Sup. Table 2-d,e.** (g) List of genes upregulated in E16 hemocytes compared to E16 glia and involved in the immune deficiency and Toll pathways. (h) Transcriptome comparison of E16 neurons and hemocytes. MA plot as in (a), the y-axis reports the log2 fold change E16 neurons/E16 hemocytes. Genes significantly upregulated in E16 neurons are shown in blue, genes significantly upregulated in E16 hemocytes in red, and non-differentially expressed genes in gray.

The comparative analysis between E16 glia and hemocytes reveals a much higher number of transcripts specifically enriched in either cell type. This is in line with the Venn diagram in **Figure 1-b**. 1945 transcripts are enriched in hemocytes (in red) and 2937 in glia (in yellow) (**Figure 2-d**). Interestingly, the hemocyte enriched transcripts only highlight one GO term related to immunity, ‘positive regulation of immune functions’, which comprises 39 genes (**Figure 2-e, Sup. Table 2-d)**. Among them are 14 members of the Toll pathway and 6 of the Immune deficiency (Imd) pathway (**Figure 2-g**). The Imd pathway is one of the conserved NF-κB (nuclear factor kappa-light-chain-enhancer of activated B cells) immune signaling pathways in insects that regulates an antibacterial defense response. The remaining GO terms include “macroautophagy” and “Golgi vesicle transport”. In the case of glia, we find glia-specific functions such as septate junction assembly (involved in BBB establishment), glial cell differentiation and neural functions such as motoneuron axon guidance (Abbott, 2005; Barres, 2008) (**Figure 2-f, Sup. Table 2-e)**.

The comparison between E16 neurons and hemocytes also shows poor similarity, with 5261 genes non differentially expressed, 3608 upregulated in neurons, 1990 in hemocytes (**Figure 2-h**).

Altogether, these data reveal that only a minority of transcripts is cell-specific and that embryonic glia and neurons are closer to each other than to hemocytes. The genes that are specifically upregulated in E16 recapitulate the specific functions of neurons and glia, while few cell-specific terms are identified in hemocytes.

### Over development, hemocytes display higher plasticity compared to neurons and glia

Cell functions adapt to changing environments during the life cycle. Typically, embryos live in autarchy, whereas larvae function as open systems, due to feeding, respiration and exposure to pathogens. Accordingly, although already functional in the embryo, we found that the differentiated cells become increasingly specialized by the larval stage.

A pairwise comparison reveals that 2500 genes are upregulated in L3 neurons compared to glia and 2279 genes are upregulated in L3 glia compared to neurons **(compare Figure 3-a and Figure 2-a**). The genes upregulated in L3 neurons highlight neuronal functions and molecular cascades linked to signaling (neuropeptide signaling pathway, regulation of neurotransmitter levels and synapse organization, **Figure 3-b, Sup. Table 3-a)**, compared to the GO terms linked to development found at E16 (**Figure 2-b**). L3 glia display the GO term septate junction assembly also found at E16, likely due to the fact that the BBB starts forming in the embryo and continues to expand as the CNS grows during the larval stages (reviewed in (Limmer et al., 2014). In addition, terms related to the differentiation of glia and gliogenesis were also found (**Figure 3-c, Sup. Table 3-b)**, in line with the extensive glial proliferation ongoing in the larva (Pereanu et al., 2005).

**Figure 3:**
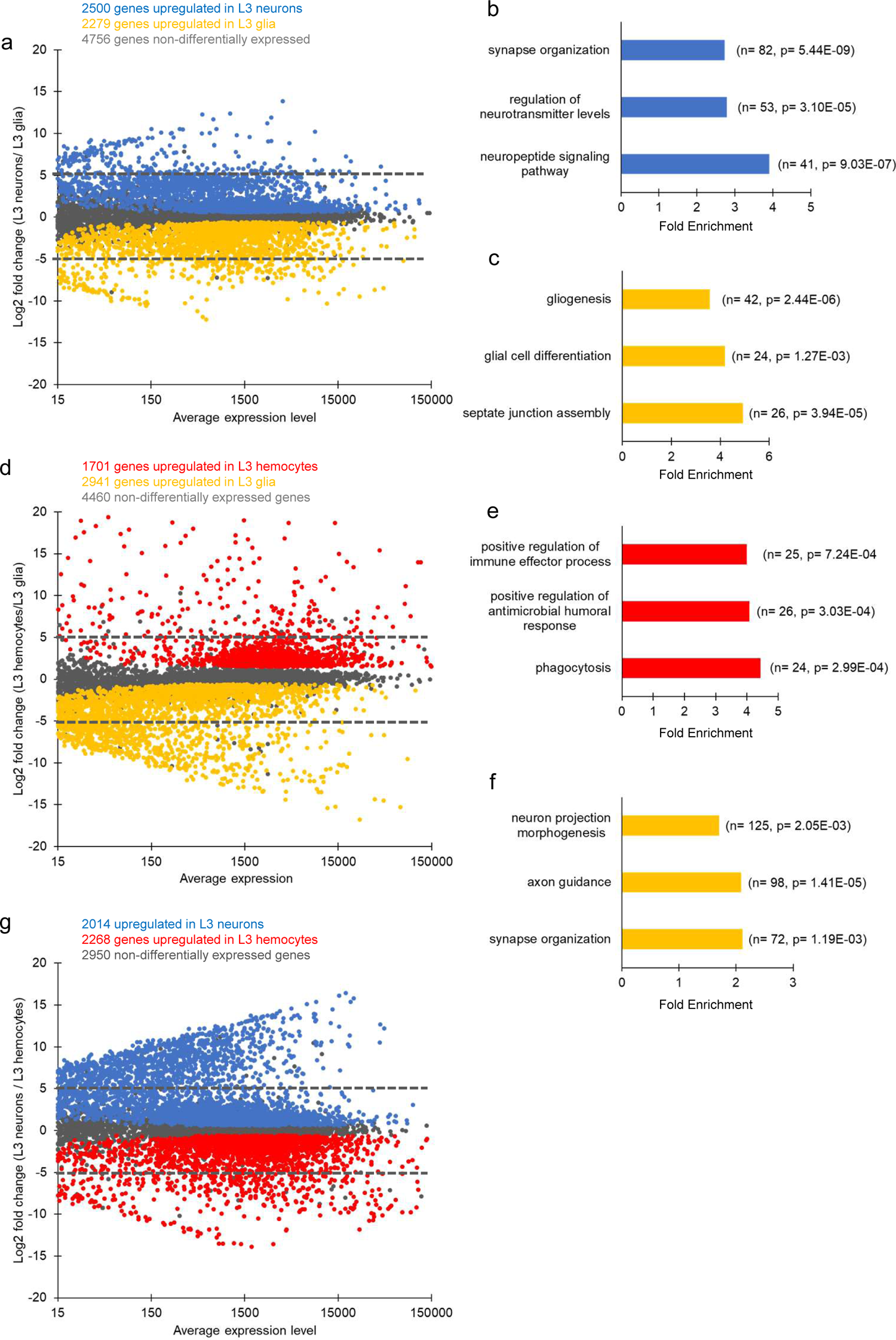
Pairwise transcriptome comparisons in the larva. (a) Transcriptome comparison of L3 neurons and glia, MA plot. The x-axis reports the average gene expression levels, the y-axis, the log2 fold change L3 neurons/ L3 glia. Genes significantly upregulated in L3 neurons are shown in blue, genes significantly upregulated in L3 glia are shown in yellow and non-differentially expressed genes are shown in gray. (b,c) Gene Ontology (GO) term enrichment analysis in L3 neurons (blue) and L3 glia (yellow). The fold enrichment for a subset of representative GO terms are displayed on the x-axis, the number of genes and the P-value of the GO term enrichment are indicated in brackets. The complete results of the GO term analysis are shown in **Sup. Table 3-a,b**. (d) Transcriptome comparison of L3 hemocytes and glia. MA plot as in (a), the y-axis reports the log2 fold change L3 hemocytes/L3 glia. Genes significantly upregulated in L3 hemocytes are shown in red, genes significantly upregulated in L3 glia are shown in yellow and non-differentially expressed genes are shown in gray. (e,f) Gene Ontology (GO) term enrichment analysis in L3 hemocytes (red) and L3 glia (yellow). The fold enrichment for a subset of representative GO terms is displayed on the x-axis, the number of genes and the P-value of the GO term enrichment are indicated in brackets. The complete results of the GO term analysis are shown in **Sup. Table 3-c,d.** (g) Transcriptome comparison of L3 neurons and hemocytes. MA plot as in (a), the y-axis reports the log2 fold change L3 neurons/L3 hemocytes. Genes significantly upregulated in L3 neurons are shown in blue, genes significantly upregulated in L3 hemocytes are shown in red, and non-differentially expressed genes are shown in gray.

We performed the pairwise comparison between L3 hemocytes and glia. Unlike their embryonic counterpart, L3 hemocytes present several terms related to immune functions such as phagocytosis, positive regulation of antimicrobial humoral response and positive regulation of immune effector process **(compare Figure 2-e and Figure 3-e, Sup. Table 3-c).** The clear shift in the hemocyte expression profile indicates a major change in the function of these cells during development. The genes upregulated in glia (2941 genes in yellow) are involved in glial functions such as synapse organization and axon guidance (**Figure 3-f, Sup. Table 3-d**), a function that has already been investigated (reviewed in (Bittern et al., 2021).

The MA plot comparing larval neurons and hemocytes shows that these cell types display the lowest level of similarity, comparable to that observed in the embryo (**Figure 3-g**).

The comparative analyses show that the divergence between neurons and glia increases over time: higher number of upregulated genes at L3 compared to E16 (Compare **Figure 3-a to Figure 2-a**,), increased fold change of upregulated genes at L3 (**Sup. Figure 2-a**). By contrast, in the case of hemocytes and glia, the number of upregulated genes as well as their median fold change value are similar in E16 and L3 (compare **Figure 2-d and Figure 3-d, Sup. Figure 2-a**).

To shed light on the hemocyte / glia similarities at L3, we identified the genes that are upregulated in L3 glia and hemocytes but not in neurons (**Figure 1-c genes in orange)**. A GO term analysis highlights genes involved in immune processes and in metabolism (**Sup. Table 4-a**). Typically, 4 genes are involved in fatty acid oxidation including CPT2, Mtpalpha, Echs1, Mfe2, CG4860 and CG17544 (**Sup. Figure 2-b, Sup. Table 4-b**), indicating the significant reliance of L3 glia and hemocytes on lipid break-down. Moreover, the upregulated genes in the L3 hemocytes and glia show several GO terms related to immune functions such as positive regulation of antimicrobial humoral response and positive regulation of immune effector process **(Sup. Figure 2-b, Sup. Table 4-a,c).**

To better understand the nature of this evolving cell behavior, we compared each cell type at E16 and L3. The MA plots show that neurons and glia evolve in a similar manner: a comparable number of genes is non-differentially expressed in neurons and in glia between E16 and L3; also, many more genes are upregulated in E16 glia and neurons, than in L3 glia and neurons (**Sup. Figure 2-c,d**). By comparison, the number of non-differentially expressed genes in E16 and L3 hemocytes is significantly lower than that detected in glia and neurons (**Sup. Figure 2-e**). This highlights the robust shift in the hemocytes’ transcriptome during development compared to the relatively less dynamic transcriptomes of neurons and glia.

Finally, and in agreement with the gene expression profiles, neurons and glia share the highest number of transcription factors at E16 (**Sup. Figure 2-f, Sup. Table 1-c,d,e**), hemocytes and glia at L3 (**Sup. Figure 2-g, Sup. Table 1-f,g,h**), further highlighting the increased commonality between hemocytes and glia at L3.

In sum, cell-specific features and specialization increase over time, especially for hemocytes, which acquire a stronger immune potential and share more similarities with glia in the larva.

### Glia and hemocytes express specific scavenger receptors

The vast majority of the hemocytes (95%) act as scavenger cells in homeostatic and challenged conditions. In the embryo, they are necessary for the elimination of apoptotic bodies and for the clearance of cellular debris. Prior to the formation of the BBB, they also contribute to this function within the developing CNS (Franc et al., 1996; Kurant et al., 2008; Manaka et al., 2004; Sonnenfeld and Jacobs, 1995b). Once glial cells form the BBB, however, hemocytes no longer have access to the CNS and by late embryogenesis the scavenging function within the nervous system is taken up by glia solely (Kurant et al., 2008). Thus, hemocytes and glia represent respectively the macrophages acting outside and inside the nervous system. The question then arises as to whether glia express the same immune pathways of hemocytes or represent a distinct class of macrophages.

A key molecular signature of macrophages is the expression of scavenger receptors (Kocks et al., 2005; Kurucz et al., 2007; Lebestky et al., 2000; Lemaitre et al., 1996; Rämet et al., 2002; Tepass et al., 1994), which we therefore used to compare hemocytes and glia. The MA plots comparing E16 (**Figure 4-a, Sup. Table 5-a)** or L3 hemocytes and glia (**Figure 4-b, Sup. Table 5-b)** allowed us to subdivide the receptors based on their relative levels of expression and fold change between the two cell types. Group 1 is specific to hemocytes, group 2 is expressed in both cell types, group 3 is specific to glia (**Figure 4-a,b group 1 in red, group 2 in black, group 3 in yellow**).

**Figure 4:**
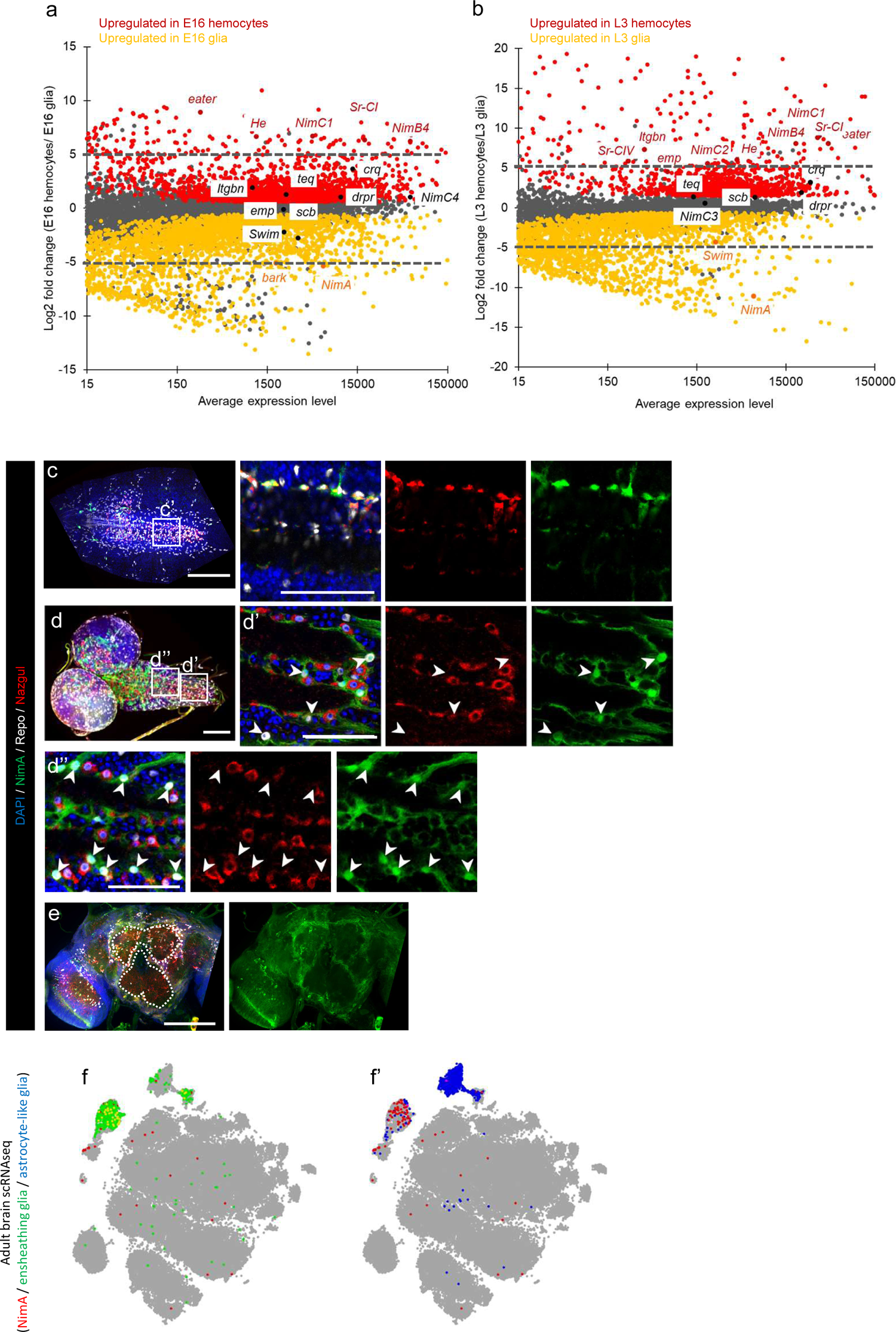
*NimA,* a novel glia-specific scavenger receptor. (a,b) MA plots as in Figure 2-c and Figure 3-c comparing the scavenger receptors expressed in embryonic and larval hemocytes and glia. In black, scavenger receptors non-differentially expressed between E16 (a) or L3 hemocytes and glia (b). In red, scavenger receptors that are upregulated in E16 (a) or L3 hemocytes (b) and in yellow, scavenger receptors that are upregulated in E16 (a) or L3 glia (b). (c,e’) Immunolabeling of *NimAGal4;UAS-eGFP* embryos (c,c’), L3 (d,d’,d’’) and adults (e) with anti GFP (green), anti-Repo (white), anti-Nazgul (red) and DAPI (blue). (c) Full stack projection, scale bar = 100 μm. Square indicates region in c’. (c’) Single section, scale bar = 50 μm. (d) Full stack projection, scale bar = 100 μm. Squares indicate regions in d’ and d’’. (d’-d’’) Single section, scale bar = 50 μm. (e) Full stack projection, scale bar = 100 μm.

Group 1 receptors are involved in the phagocytosis of Gram-positive and Gram-negative bacteria (Pearson et al., 1995; Rämet et al., 2001), matching the hemocyte-specific function of pathogen phagocytosis. They include the well-studied genes *eater, NimC1, NimC2* and *Scavenger receptor class C, type I (Sr-CI)*. *eater, NimC1* and *NimC2* contain multiple epidermal growth factor (EGF) like repeats called NIM repeats (Kurucz et al., 2007). SR-CI and SR-CIV belong to the *Drosophila* SR-C class of scavenger receptors, SR-CI contains several known motifs, including two complement control protein domains, a somatomedin B domain and a Meprin, A-5 protein, and receptor protein-tyrosine phosphatase Mu domain (Rämet et al., 2001).

Group 2 receptors include *NimC4, draper (drpr) and croquemort (crq*), which was believed to be only expressed in hemocytes (Franc et al., 1996) and belongs to the SR-B class of scavenger receptors homologous to the mammalian CD36 (Franc et al., 1999). Although *crq* is expressed at higher levels in hemocytes, it is still expressed in glia at relatively high levels and cannot be considered hemocyte-specific **(Sup. Table 5-a,b)**. These receptors are involved in the phagocytosis of apoptotic bodies and debris, in line with the fact that Drpr and NimC4 act in hemocytes and in glia to promote the clearance of apoptotic cells and axonal debris (Kurant et al., 2008; MacDonald et al., 2006; Manaka et al., 2004; Melcarne et al., 2019; Roddie et al., 2019).

Group 3 includes three genes: *bark beetle* (*bark*) encodes a transmembrane scavenger receptor-like protein important for the establishment of the BBB (Hildebrandt et al., 2015; Parker and Auld, 2006; Stork et al., 2008). *Secreted Wg-interacting molecule* (*Swim*) contains a somatomedin B domain similar to Sr-CI, however it has not been investigated for its role in phagocytosis. NimA is a Drpr-like receptor containing one NIM domain followed by two EGF repeats and one Emilin domain (**Sup. Figure 3-a**, Callebaut et al., 2003). Its *C. elegans* orthologs are involved in the phagocytosis of apoptotic cells (Kurucz et al., 2007; Mangahas and Zhou, 2005).

The expression profile of the receptors evolves with time. The number of scavenger receptors specific to hemocytes and their expression levels increases in the larva (4 at E16, 8 at L3, shown in red, **Figure 4-a,b**). In parallel, the number of receptors non-differentially expressed between hemocytes and glia decreases in the larva (8 at E16, 5 at L3, shown in black). Moreover, within the same cell type, the expression levels of some receptors change significantly, *eater* being highly upregulated in L3, *NimC4* at E16.

To validate the expression of these genes, we performed quantitative reverse transcription PCR (RT-qPCR) assays on L3 glia and hemocytes. We tested and confirmed the purity of the sorted cell populations by quantifying the expression levels of *Hemolectin* (*Hml)* and *reverse polarity* (*repo)*, as markers of hemocytes and glia, respectively (**Sup. Figure 3-b,c**). *NimC1* and *eater* are significantly upregulated in L3 hemocytes compared to glia **(Sup. Figure 3-d,e)**, *drpr* shows comparable expression levels **(Sup. Figure 3-f);** *crq* seems to be more expressed in hemocytes however the difference is not statistically significant **(Sup. Figure 3-g)**, similarly to the transcriptome results. *NimA* is only found in glia **(Sup. Figure 3-h)**. The finding that some receptors are expressed in glia and hemocytes, whereas others are cell-specific recapitulates their common function of phagocytosing apoptotic cells and debris and at the same time emphasizes the role of hemocytes in host defense against pathogens. In addition, the higher levels of the scavenger receptors in L3 than in E16 hemocytes prove an increased functional specificity of hemocytes in the larva.

Given the high specificity and levels of expression of *NimA*, we performed an *in situ* hybridization RNAscope assay on wild type (*WT)* L3 CNS and found very specific labeling at the position of the neuropile (**Sup. Figure 3-i,i’**). We complemented this analysis by labeling *NimA-Gal4; UAS-eGFP* E16, L3 and adults with Repo, a pan-glial marker and Nazgul, a NADP-retinol dehydrogenase specifically expressed in embryonic astrocyte and ensheathing glia, the two cell types that constitute the neuropile glia (Peco et al., 2016; Ryglewski et al., 2017) (**Figure 4-c-e’**). Nazgul and NimA nicely colocalize in the embryonic glial cells (**Figure 4-c,c’**). By contrast, NimA and Nazgul are found in larval glial cells that surround the neuropile and are in close proximity, suggesting that at that stage Nazgul is expressed in the astrocyte glia that invade the neuropile, while *NimA* in expressed in the ensheathing glia as well as in the wrapping glia that surround the peripheral nerves (**Figure 4-d,d’**). In the adult CNS, *NimA* is also found in glial cells surrounding the neuropile and, based on single-cell RNA seq data, *NimA* is expressed in adult ensheathing glia but not in astrocyte-like glia (**Figure 4-f,f’**) (Davie et al., 2018). Thus, NimA represents a novel glia-specific scavenger receptor expressed in a subset of glia cells throughout the life of the fly.

In sum, hemocytes have a strong commitment to immune functions in L3 and each phagocytic cell type expresses common as well as specific receptors, such as the glia-specific *NimA* gene and the hemocyte-specific *NimC1* and *eater* genes. This highlights a cell-specific phagocytic potential that differentiates glia from the macrophages outside the CNS.

### Stage-specific transcriptional signatures

The transition from the embryo to the larva involves massive physiological changes that affect the organism as a whole. This likely explains why the transcriptomes group by stage more than cell types (**Figure 1-a**). To get more mechanistic insights on the changes associated with time, we assessed whether a stage-specific molecular signature is common to all cell types. We combined all cells deriving from E16 and compared their transcriptomes to those of cells deriving from L3.

The MA plot comparing E16 to L3 transcriptomes suggests that embryonic cells are overall closer to each other than larval cells (**Figure 5-a**), in line with the dendrogram in **Figure 1-a**. This does hint to a stage-specific signature common to all cells. The transcripts enriched at E16 compared to L3 are involved in cytoplasmic translation and cuticle development (**Figure 5-b, Sup. Table 6-a**). Genes included in the GO term cytoplasmic translation code for ribosomal proteins and translation machinery, such as translation initiation factors. Ribosomal protein transcripts and ribosomes are known to be maternally deposited and highly abundant in early embryos (Qin et al., 2007), but new ribosomes are built by the end of embryogenesis, in line with the transcriptomic data. The ‘cuticle development genes’, include 42 genes coding for Cuticular proteins, 21 genes coding for members of the Tweedle family and 7 genes coding for CPR cuticle protein family (**Figure 5-c, Sup. Table 6-b**). These proteins are predicted to be present in the extracellular matrix (ECM) (Cornman, 2009; Karouzou et al., 2007; Naba et al., 2016; Zuber et al., 2020), which implies a role of all three cell types in the deposition of ECM and cuticle at embryonic stage, a feature that has already been described for the hemocytes (Brown, 2011; Martinek et al., 2008) and is in accordance with the important role of the ECM in organogenesis.

**Figure 5:**
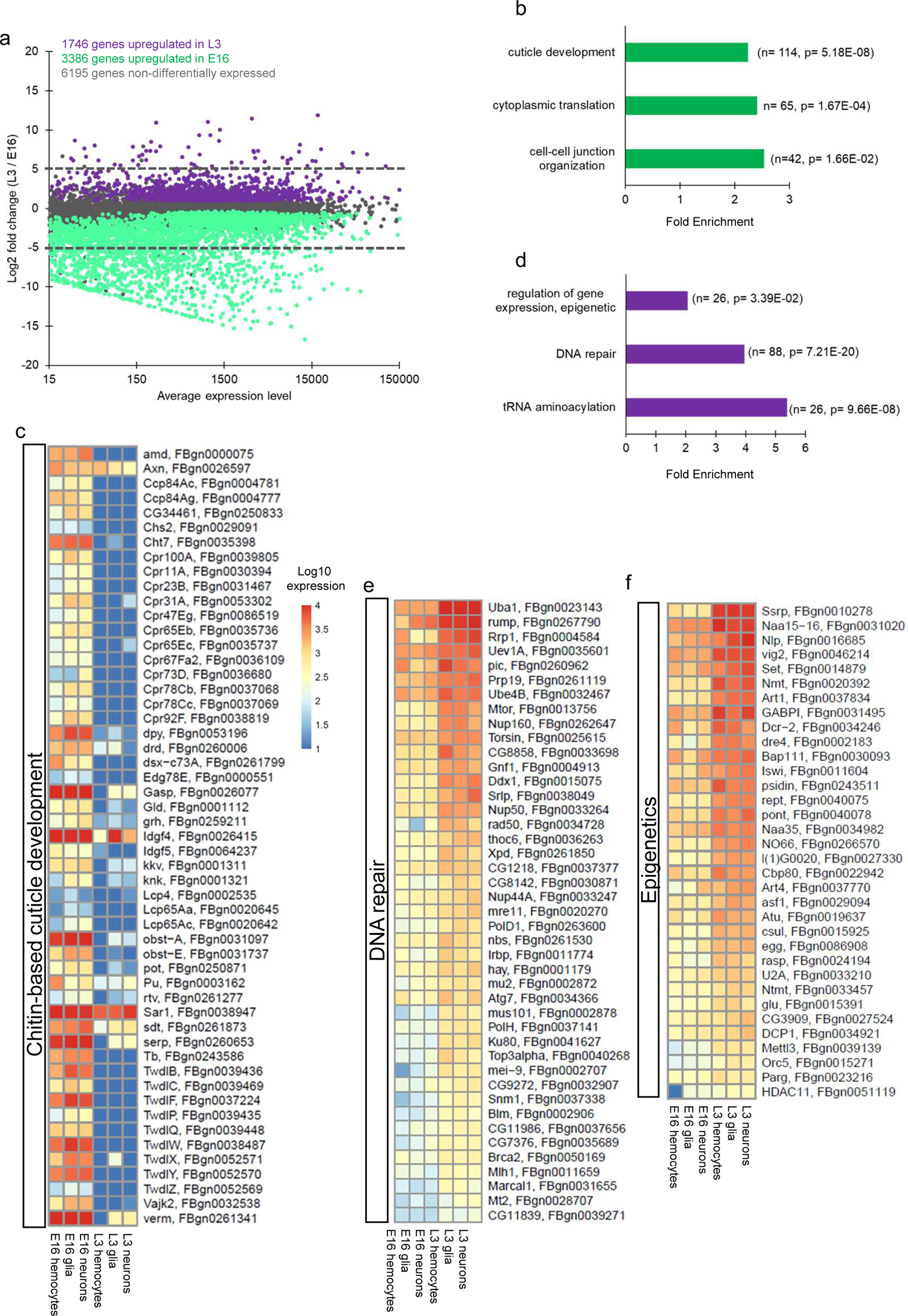
Pairwise transcriptome comparison between E16 and L3 cells. (a) Transcriptome comparison of L3 vs. E16 cells, MA plot. The x-axis reports the average gene expression levels, the y-axis, the log2 fold change L3 cells/E16 cells. Genes significantly upregulated in E16 are shown in green, genes significantly upregulated in L3 are shown in purple and non-differentially expressed genes are shown in gray. (b,d) GO term enrichment analysis in L3 (purple) and E16 (green). The fold enrichment for a subset of representative GO terms is displayed on the x-axis, the number of genes and the P-value of the GO term enrichment are indicated in brackets. The complete results of the GO term analysis are shown in **Sup. Table 6-a,c.** (c,e,f) Heatmaps showing the log10 gene expression of genes upregulated in E16 cells compared to L3 cells and involved in Chitin-based cuticle development (c), as well as that of genes upregulated in L3 cells compared to E16 cells and involved in DNA repair (e) and epigenetics (f). Scale shows low expression (blue) to high expression (red). Values for heatmaps shown in **Sup. Table 6-b,d,e**.

Fewer genes are upregulated in L3 compared to E16 (**Figure 5-a**) and many of them belong GO terms related to tRNA aminoacylation, DNA repair and epigenetics (**Figure 5-d, Sup. Table 6-c**). The enrichment of tRNA aminoacylation related transcripts hints to increased translation, which could be due to the fact that, unlike embryos, larvae take up amino acids and nutrients through feeding and are thus able to aminoacylate tRNAs, as lack of amino acids and starvation has been shown to cause significant decreases in protein synthesis (Deliu et al., 2017).

We were puzzled by the enrichment of transcripts involved in DNA repair. Among them are *rad50*, *Nijmegen breakage syndrome (nbs)* and meiotic recombination 11 (*mre11*), members of the *MRN* complex, as well as *Telomere fusion (Tefu*), the ortholog of *Ataxia Telangiectasia Mutated (ATM)* (**Figure 5-e, Sup. Table 6-d)**. These genes have been implicated in DSB repair (Wang et al., 2014), in DNA replication restart after replication stress (Gatei et al., 2014) and in preventing telomere fusion as well as chromosome breakage in *Drosophila* (Ciapponi et al., 2004, 2006). To validate the stage-specific enrichment of these transcripts, we performed qPCR assays on RNA extracted from whole E16 and L3 animals: *rad50* (**Sup. Figure 4-a)** and *mre11* (**Sup. Figure 4-b)** are upregulated in L3 and *nbs* tends to be upregulated **(Sup. Figure 4-c)**. Based on the fact that double strand breaks trigger the phosphorylation of the histone H2AX, a variant of the H2A protein family (Redon et al., 2002), we hypothesized that the increase in DNA repair genes is due to an increase in DSBs. Therefore, we estimated the relative levels of DSB using the phosphorylated H2A.v (H2A.v-P) *Drosophila* marker, the equivalent of gamma-H2AX in mammals. When DNA damage forms double strand breaks, H2A.v is phosphorylated by kinases such as ATM and the phosphorylated protein, gamma-H2AX, is the first step in recruiting and localizing DNA repair proteins. Western blot assays on histone extracts show that the levels of H2A.v-P decrease in L3 compared to E16 **(Sup. Figure 4-d,e)**, contrary to our initial hypothesis.

A recent study showed that the yeast equivalent of the MRN complex (MRX) is involved in controlling gene expression levels through modulating chromatin interaction (Forey et al., 2021). Moreover, MRN components interact with Heterochromatin Protein 1 (HP1), which is involved in gene repression through heterochromatin formation (James and Elgin, 1986; James et al., 1989), and they regulate HP1 expression and stability (Bosso et al., 2019). This, in addition to the finding of GO terms related to epigenetics “negative regulation of gene expression, epigenetic”, “chromatin remodeling” and “histone modification” in the genes upregulated in L3 compared to E16 (**Figure 5-f, Sup. Table 6-e**) led us to ask whether Rad50 might be involved in chromatin conformation and in the regulation of gene expression.

Chromatin states are defined by a highly complex epigenetic code comprised of histone post-translational modifications. Histone methylation is complex in that residues may be mono-, di-, or trimethylated, and these marks may activate or repress gene transcription. Di- and trimethylation of H3K4, for instance, strongly correlate with active gene transcription, whereas H3K9 and H3K27 methylation are mostly associated to a closed heterochromatin state and therefore to transcription repression (Aranda et al., 2015; Conway et al., 2015; Peters et al., 2003; Rea et al., 2000; Santos-Rosa et al., 2002). To determine whether Rad50 is required for chromatin conformation and repression of gene expression in L3 larvae, we labeled polytene chromosomes of wild-type or *rad50Δ5.1* mutant larvae with antibodies against H3K4me3, H3K9me2 or H3K9me3. The absence of Rad50 causes a significant increase of H3K4me3 and a decrease of H3K9me2/me3 epigenetic marks (**Figure 6-a,b**). Polytene chromosomes are characterized by the alternation of dense (heterochromatic) and less dense (euchromatic) bands, a reflection of the chromatin condensation state. Interestingly, we observed an alteration of the H3K4me3 band signal profile accompanied by almost complete absence of H3K9me3 in *rad50Δ5.1* polytene chromosomes compared to *WT* (**Figure 6-c,d).** While the repression marks H3K9me2/me3 show a clear decrease in *rad50Δ5.1,* the H3K27me3 signal increases significantly in *rad50Δ5.1* (**Figure 6-e,f**). Due to the link between H3K27me3 and DNA damage shown by studies highlighting an increase in H3K27me3 at DNA DSB sites (Basenko et al., 2015; O’Hagan et al., 2008, 2011) and the role of Rad50 in DNA damage, we asked whether the increase in H3K27me3 is simply due to increased DSBs. Comparing H3K27me3 and H2A.v-P distribution shows a clear lack of colocalization of the two marks thus highlighting that the increase in H3K27me3 in independent from the increase of DSBs (**Figure 6-e-g**).

**Figure 6:**
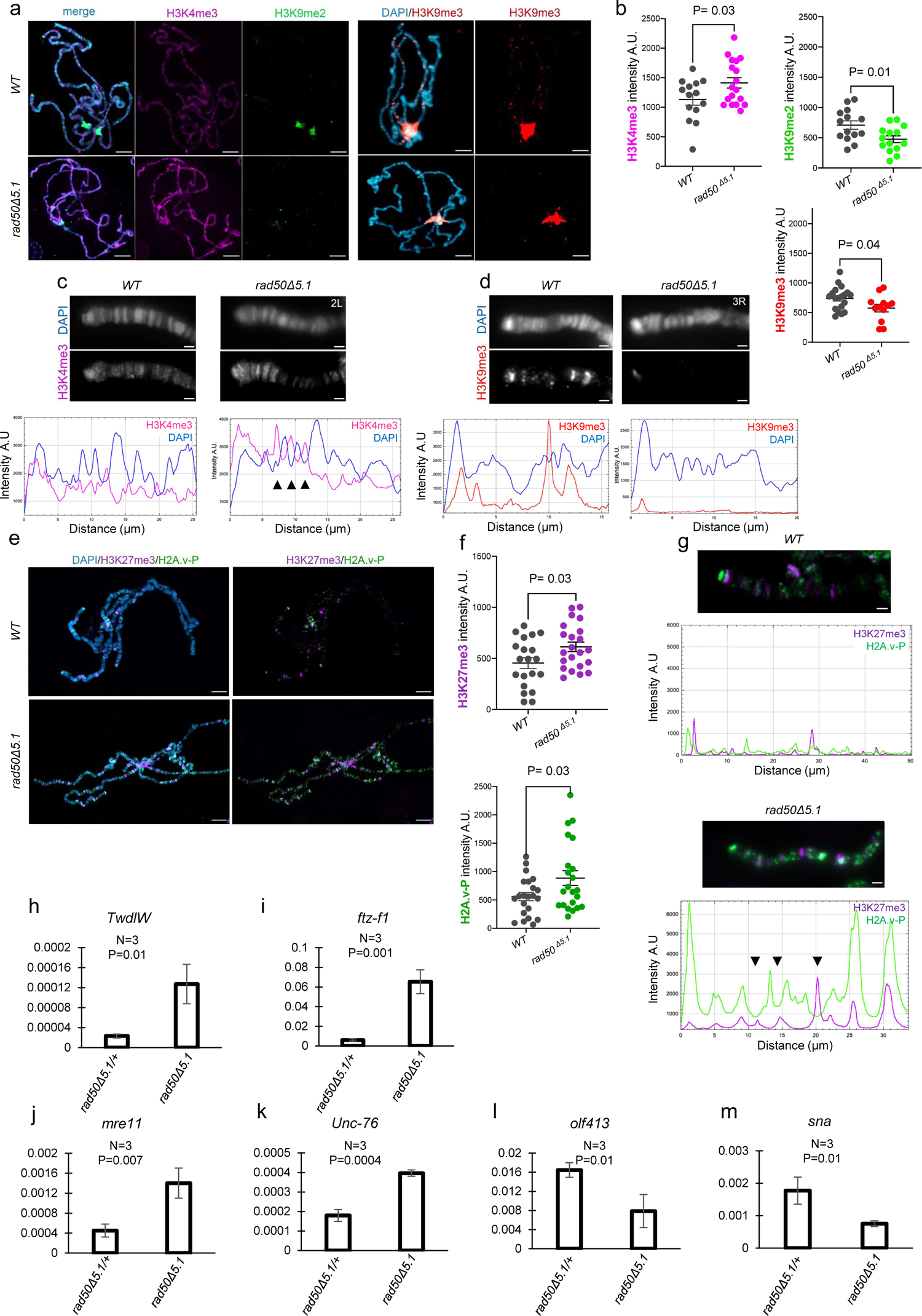
Impact of *rad50* on polytene chromosomes and on gene expression. (a) Representative polytene chromosomes immunolabeled with anti-H3K4me3 (magenta) and anti-H3K9me2 (green), or with anti-H3K9me3 (red), from *WT* or *rad50Δ5.1* homozygous mutants L3. Chromosomes were counter-stained with DAPI (blue). Scale bar = 20 μm. (b) Fluorescence intensity quantification of polytene chromosomes, labeled as in (a), shows an increase of the euchromatic marker H3K4me3 and a decrease of the heterochromatic markers H3K9me2 H3K9m3, in *rad50Δ5.1* mutants compared to *WT*. Each dot represents a single polytene chromosome (n >/= 13). Arbitrary Unit (A.U.). Error bars represent SEM. P = p-value calculated using Welch parametric t-test. (c) Representative intensity profiles of H3K4me3 and DAPI signals over a segment at the tip of chromosome 2L, showing an alteration of the H3K4me3 signal in *rad50Δ5.1* mutant compared to the *WT*; arrowheads denote regions of strong difference of H3K4me3 in *rad50Δ5.1* compared to *WT*. y-axis: fluorescence intensity; x-axis: distance from the tip of the chromosome (μm). Scale bar = 10 μm. (d) Representative intensity profiles of H3K9me3 and DAPI signals over a segment at the tip of chromosomes 3R showing absence of H3K9me3 signal in *rad50Δ5.1* compared to *WT*. y-axis: fluorescence intensity; x-axis: distance from the tip of the chromosome (μm). Scale bar = 10 μm. (e) Representative polytene chromosomes immunolabeled with anti-H3K27me3 (purple) and anti-H2A.v-P (green), from *WT* or *rad50Δ5.1* homozygous larvae. Chromosomes were counter-stained with DAPI (blue). Scale bar = 20 μm. (f) Fluorescence intensity quantification showing increase of both H3k27me3 and H2A.v-P signals in *rad50Δ5.1* mutants compared to *WT*. Each dot represents a single polytene chromosome (n >/= 15). Error bars represent SEM. P = p-value calculated using Welch parametric t-test. (g) Representative intensity profiles of H3K27me3 and H2A.v-P signals over a segment at the tip of the 2R chromosome. Note that the increase of H2A.v-P peaks in the *rad50Δ5.1* mutant is not directly correlated with that of H3K27me3 (arrowhead). y-axis: fluorescence intensity; x-axis: distance from the tip of the chromosome (μm). Scale bar = 10 μm. (e-j) RT-qPCR assays on *rad50Δ5.1 /+ and rad50Δ5.1* larvae. The levels of *TwdlW* (h), *ftz-f1* (i), *mre11* (j), *Unc-76* (k), *olf413* (l) and *sna* (m) transcripts were quantified and the relative expression levels were calculated based on the expression of *Act5C* and *RP49*. List of primers shown in **Sup. Table 9,** n = number of biological replicates, P = p-value calculated using Student’s t-test and indicated on graphs, error bars represent the standard deviation.

The changes in the H3K4me3 and H3K27me3 profiles and the strong reduction of H3K9me3, observed in *rad50Δ5.1* polytene chromosomes, suggest that Rad50 depletion affects chromatin organization and hence gene expression regulation. These findings, together with the published data, underline a novel role of the DNA damage factors, in particular Rad50, in regulating chromatin states.

To understand whether Rad50 depletion and its effect on chromatin state also impinge on gene expression, we analyzed several transcripts that show high expression in E16 but low expression in L3 in all cell types in the *WT*. Assessing the expression of these genes in L3 *WT* compared to *rad50Δ5* showed significant upregulation of four genes upon Rad50 depletion including the *TweedleW* (*TwdlW*), *ftz transcription factor 1* (*ftz-f1*)*, Uncoordinated 76 (Unc-76)* and the member of ATM *Mre11* (**Figure 6-h,k**), while *olf413* and *snail* (*sna*) show significant decrease of expression levels (**Figure 6-l,m**). Altogether, these and the published data highlight a novel role of the DNA damage factors, and specifically Rad50, in transcriptional regulation.

In sum, our data reveal that different developmental stages are associated with specific molecular signatures, regardless of the cell type, including chromatin organization.

### Metabolic signatures of embryonic and larval neurons, glia and hemocytes

A careful comparison of the embryonic and larval cells also revealed distinct metabolic states. Compared to their larval counterpart, E16 hemocytes express more glycolytic enzymes such as *lactate dehydrogenase (ldh)*, essential for the conversion of pyruvate to lactate (Cattenoz et.al., 2020, **Figure 7**). This implies the activation of a Warburg-like glycolytic state and the conversion of pyruvate to lactate. E16 hemocytes are also lipogenic as they highly express *de novo* the lipogenic enzyme, *Acetyl-CoA carboxylase (ACC)* and enzymes of the triacylglycerol biosynthetic pathway (TAG) that include *Fatty acid synthase (FAS)*, *Glycerol-3-phosphate acytransferase* (*Gpat), 1-Acylglycerol-3-phosphate O-acyltransferase* (*Agpat), Lipin (Lpin)* and *Diacylglycerol O-acyltransferase (DGAT)*. By contrast, L3 hemocytes are more oxidative, they most likely internalize and metabolize lipids through the beta-oxidation pathway to generate acetyl CoA and drive the tricarboxylic acid (TCA) cycle. This is supported by the enrichment of transcripts encoding lipid-scavenging receptors and by the poor expression of genes involved in the TAG pathway (**Figure 7, Sup. Table 7)**. Moreover, L3 hemocytes do not rely on glycolysis to drive the TCA cycle, as the genes involved in glycolysis, mainly *phosphofructokinase* and *pyruvate dehydrogenase* (*PDH*), of which *PDH* controls the rate-limiting conversion of pyruvate to acetyl CoA and critically controls TCA activity, are not upregulated in those cells. Finally, the poor expression of *phosphoenolpyruvate carboxykinase* (*PEPCK*) and *fructose 1,6-bisphosphatase (FBPase),* as well as a significant enrichment of the *Glut1* sugar transporter transcripts suggest the absence of gluconeogenesis and the active uptake of glucose in the larva. Because the glycolytic genes downstream of *glucose 6, phosphate (G6P)* are poorly expressed while those of the pentose phosphate pathway (PPP) are highly expressed in the L3 hemocytes, the internalized glucose could be used to generate pentose sugars for ribonucleotide synthesis and redox homeostasis through the generation of NADPH. Hence, apart from the glycolytic pathway, E16 and L3 hemocytes display anabolic (gluconeogenic, lipogenic) and catabolic (oxidative, lipolytic) signatures, respectively. Comparative analyses revealed interesting commonalities in a few metabolic pathways, in addition to metabolic changes that are cell-specific, which projects underlying differences in the metabolic states (**Figure 7**). The one feature observed across all cell types is a clear metabolic shift in the key glycolytic enzyme *ldh*, which is highly expressed in the embryo (**Figure 7, Sup. Table 7**). This implies a global activation of a Warburg like state specifically in the embryo.

**Figure 7:**
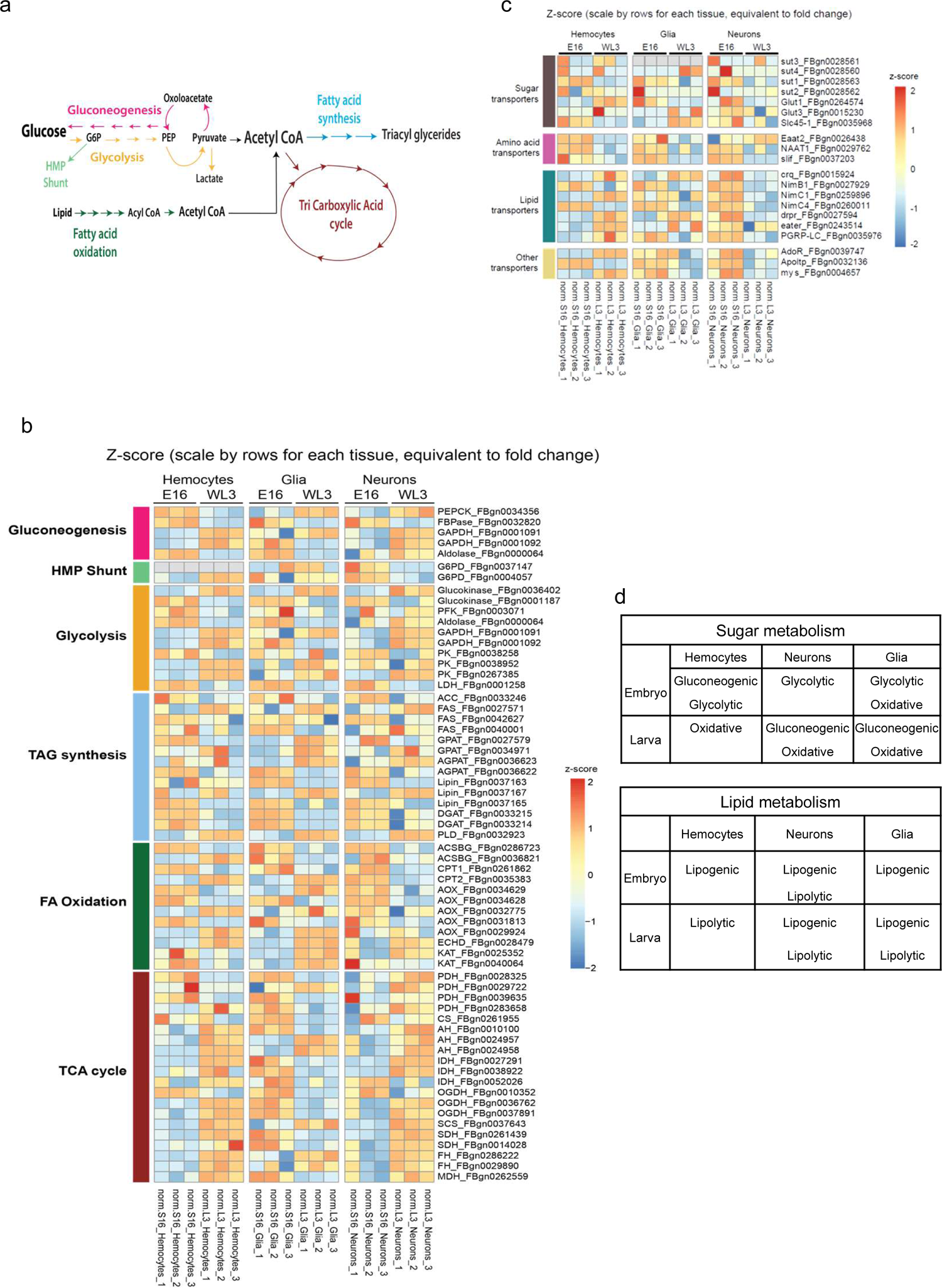
Comparison of the metabolic states in hemocytes, glia and neurons at E16 and L3. (a) Schematic showing the main steps of sugar and lipid metabolic pathways. (b,c) Heatmaps showing the z-score of genes involved in the different metabolic processes (b) and transporters (c) in hemocytes, glia and neurons at E16 and L3. Scale shows low expression (blue) to high expression (red). Values are shown in **Sup. Table 7.** (d) Recapitulative table highlighting the main processes for sugar and lipid metabolism in hemocytes, neurons and glia from E16 and L3.

The transcripts of the glycolytic pathway genes are enriched in E16 glia and neurons: *Glucokinase (Hexokinase-C, FBgn0001187), Aldolase* and *Glyceraldehyde 3-phosphate dehydrogenase (Gapdh)* show prominent expression in glia, while only *Glucokinase* and *pyruvate kinase* are expressed at high levels in the neurons. The transcripts coding for the *Glucose 6-phosphate dehydrogenase (G6PD)*, a rate-limiting enzyme of the PPP pathway, is also strongly enriched in E16 glia and neurons. This implies that these cells most likely use glucose for driving the PPP pathway and glycolysis. Neither E16 glia nor E16 neurons are gluconeogenic, as the *PEPCK* gene is poorly expressed at that stage. We therefore assessed the underlying source of glucose driving glycolysis and evaluated the capacity for glucose uptake via transporters as a predictable source. This revealed a strong enrichment of transcripts coding for sugar transporters like *sut1, sut3, sut4* and specifically glucose transporters *Glut1* and *Glut3*. L3 glia and neurons seem both gluconeogenic and glycolytic. Thus, E16 glia and neurons rely on extracellular sugar uptake, sustained glycolysis and PPP while L3 glia and neurons utilize *de novo* gluconeogenesis to fuel glycolysis.

The fate of glucose, post its breakdown to pyruvate via glycolysis, apart from driving lactate, can be diverted to the TCA cycle (oxidative). We did observe a significant upregulation the TCA enzymes in E16 glia compared to their larval counterpart. For instance, *Citrate synthase*, *Aconitase*, *Isocitrate dehydrogenase (IDH), oxoglutarate dehydrogenase (Ogdh), succinate dehydrogenase (SDH)* and *malate dehydrogenase* (*MDH*) are highly expressed in the embryo. By contrast, only a subset of TCA enzymes (*Aconitase, succinyl CoA synthetase (SCS)*, *Fumarate hydratase* (*FH))* show prominent expression in L3 glia. Thus, glia are both glycolytic and oxidative in the embryo and transition to being primarily oxidative in the larva. By contrast, neurons undergo a clear metabolic shift from glycolytic in the embryo to oxidative in the larva, as L3 neurons display high expression of most TCA enzymes, including *PDH.* Both E16 glia and neurons show significant enrichment of lipogenic transcripts. The key enzyme, *ACC,* and TAG biosynthetic enzymes (*FAS, Gpat, Agpat, Lpin*) are highly expressed. Interestingly, the expression of enzymes involved in lipid break down, such as *Acyl-CoA oxidase (AOX)*, the first enzyme of the beta-oxidation system, and *Carnitine O-palmitoyltransferase (CPT1* and *CPT2)*, which transfers fatty acids into the mitochondrion for β-oxidation, is also high in E16 neurons but not glia. Thus, L3 glia continues to show elevated expression of a subset of TAG biosynthetic enzymes (*FAS, Gpat, Agpat, Lpin*). Additionally, L3 glia show high expression of the fatty acid breakdown pathway. Taken together, these data suggest that glia are lipogenic both in embryo and larvae but activate lipolytic genes specifically in the larval stage. Neurons, on the other hand, sustain a lipogenic and lipolytic state in the embryo but in the larva, they are only lipogenic.

## Discussion

The definition of cell types makes the object of intense investigation for the potential impact on medical science and in particular on regenerative medicine. We here identify the molecular signatures that characterize *Drosophila* neurons, glia and hemocytes, as representative examples of related and unrelated cell types. We show that the molecular identity of a cell type results from the combined action of intrinsic and extrinsic cues and we establish the impact of origin, position, metabolic state, function and time. We identify cell-specific as well as stage-specific signatures and we show that cells maintain their identity during development while keeping their ability to accommodate to the evolving needs of the organism.

### Cell types are defined by a relatively low number of specific transcripts

Several technologies allow the genome-wide characterization of cells at the molecular level. Bulk transcriptomes provide an adapted tool to extrapolate the most significant and specific features that identify a cell type as a whole, due to the sequencing depth and to the average measurement of gene expression across a population of cells. Using this approach, we show that the majority of the genes are not differentially expressed between neurons, glia and hemocytes, both in the embryo and in the larva. Such high number of similar genes cannot be due to common origin/function because neurons and hemocytes share neither. This data nevertheless goes along with a study showing that 48.8% of all Flybase genes are non-differentially expressed between all stages (Daines et al., 2011). No more than 2000 genes are upregulated in each cell type, hence, a relatively low number of genes is sufficient to define a cell type.

In average, the transcripts enriched in a specific cell type are expressed at high levels, suggesting that they are under the control of signaling pathway rather than being expressed constitutively and at low levels. Thus, cell types are defined by a combination of cell-specific transcript enrichment and high levels of expression. Of note, even amongst the common transcripts we find a similar distinction: the genes that have a house keeping function shared by all cell types are expressed at lower levels than those that are associated with more specific functions. For instance, the comparison of neurons and glia shows that the genes involved in DNA replication display a number of reads < 300 while those related to axons (which are common to neurons and glia but not to hemocytes) are expressed at higher levels (> 3000 reads).

In sum, cell types are defined by few genes expressed at rhigh levels.

### Transcriptional landscapes and the impact of time

The establishment of a cell type involves the stable expression of specific morphological and functional features, even though cells may lose some traits or assume others over time. We hence expected each cell type to group together throughout development. Instead, the developmental stage represents the first factor of homogeneity, more than the cell type: embryonic hemocytes are closer to embryonic glia and neurons than to larval hemocytes. Thus, although differentiated cells accomplish the same function throughout the life of the organism, time affects their state.

Amongst the stage-specific genes, ECM related GO terms are upregulated in E16 compared to L3 cells. In line with the fact that the ECM is a key player in morphogenesis and organogenesis, different types of embryonic cells likely express large quantities of ECM transcripts to prepare for the larval life and the immense growth happening during this stage. L3 cells show an enrichment of transcripts involved in DNA damage and repair. In particular, the three components of the MRN complex (Rad50, Mre-11 and NBS) and Tefu, that respond to DNA DSB and are involved in replication fork dynamics, telomere maintenance and even response to viral infection (Ciapponi et al., 2004; Syed and Tainer, 2018). The increased expression of these genes is seemingly not due to DSB since there are more DBS markers in E16 than in L3. Our results suggest that Rad50 functions beyond DNA repair and controls gene expression through the modulation of chromatin state, in line with recent finding showing that the yeast ortholog represses the loci near its binding site (Forey et al., 2021). Future studies will address whether the changes in both euchromatic and heterochromatic marks observed in the mutant larval salivary glands are due to late, indirect, effects or reflect a novel regulatory mechanism.

The RNAseq data suggest that cells are able to sense time, a process that at least in the case of neurons is disconnected from the cell cycle and from cell division, as these cells do not undergo mitosis. The upregulation of genes involved in histone modification and chromatin remodeling in L3 suggests a stage-specific epigenetic control of the transcriptional landscape.

Finally, the analysis of the transcription factors at the two stages suggests that the three cell types are endowed with distinct degrees of plasticity. The E16 and the L3 hemocytes share many fewer transcription factors (n=25, **Sup. Table 1-i,j in red**) than those that are common to the three embryonic cell types (n=129, **Sup. Table 1-a).** This behavior is also found in glia (n=17, **Sup. Table 7-k,l in yellow**) but not in neurons (n= 124, **Sup. Table 1-m,n in blue**), which share more than 50% of upregulated transcription factors at E16 and L3. The transcriptome of neurons as well as the transcription factors shared at E16 and L3 suggest that neurons are a relatively stable population compared to hemocytes. In the case of glia, though comparing their transcriptome at E16 and L3 shows similar behavior to neurons, the fact that they share few transcription factors at E16 and L3 suggests an intermediate level of stability. It is tempting to speculate that the neural-immune phenotype of glial cells and the strong reliance of hemocytes on the environment could account for the plastic behavior of these cell types. An increased number of glial and hemocytes subtypes in larval stages could also account for such behavior. Future single cell analyses on the different cell populations at different stages will help distinguishing between these possibilities.

### Hemocytes and glia, macrophages outside and within the nervous system

The origin and position of neurons and glia have a significant impact on their transcriptional landscapes, as shown by the large number of common molecular features. A substantial fraction of the shared GO terms concern processes related to axons, synapses, and neural processes. In contrast, neurons and hemocytes share many fewer transcripts (5,9% at E16, 3,8% at L3), with no GO terms related to cell-specific processes (**Sup. Table 8-a,b**). The situation is different for the glia/hemocyte comparison. E16 glia and hemocytes share the same very low degree of similarity as that observed between neurons and hemocytes (5,7%). In addition, genes upregulated in E16 hemocytes do not seem to be involved in hemocyte-specific functions. This is likely due to the fact that the major function of the E16 hemocytes is to secrete ECM components and to phagocytose apoptotic cells, both of which are also done by glia. By contrast, L3 hemocytes and glia share 18,3% of enriched transcripts, some of which are related to immunity. The presence of hemocyte-specific GO in the larva indicates a real developmental switch (see also Cattenoz et al., 2020) and a strong specialization of the hemocytes, with more refined groups of genes being upregulated and a strong immune component.

Glia are not just macrophages that are also able to perform neural functions. Much like microglia, the vertebrate myeloid cells that invade the nervous system during development (Neumann et al., 2009; Wu et al., 2015) (DePaula-Silva et al., 2019; Hickman et al., 2013; Li and Barres, 2017; Ritzel et al., 2015), fly glia express very specific scavenging features compared to the immune cells located outside the nervous system. Typically, *NimA* is not expressed in hemocytes but only in a subset of glia. Pursuing the role of *NimA* will help deciphering the mode of action of glia in the immune response, which so far has relied only on the study of Draper and NimC4, the latter being only expressed in embryonic glia. NimA is likely not the only additional scavenger receptor possibly involved in the phagocytosis of debris and apoptotic bodies. A recent study identified the role of NimB4 in apoptotic bodies phagocytosis in glia and in hemocytes in the larva (Petrignani et a., 2021), even though *NimB4* expression in hemocytes is at least 70-fold higher than in glia (our data).

The RNAseq data highlight an intermediate neuro-immune phenotype of fly glia. We propose that simple organisms display relatively few cell types with multiple potentials whereas more complex organisms need a more refined division of labor accompanied by the appearance of dedicated cell types, such as the microglia in vertebrates. Interestingly, the comparative analysis of the transcription factor reveals the upregulation of the Nuclear factor of activated T-cells (NFAT) ortholog in L3 glia and hemocytes compared to neurons **(Sup. Table 1-g in gray)**. While the *Drosophila* gene has not yet been explored in the context in immunity, NFAT is a member of a Calcium dependent protein family that plays an important role in the immune response in lymphoid and myeloid vertebrate lineages, including microglia (reviewed in Müller and Rao, 2010).

In sum, hemocytes become more specialized as the animal hatches. Fly glia are at the interface between the nervous and the immune systems: the neural origin shared with neurons is already apparent in the embryo, while the immune function shared with hemocytes becomes apparent in the larva. Given the importance of the immune resident cells of the CNS in development and homeostasis, fly glia represent an interesting tool to unravel the role of myeloid-like cells in diseases as severe and diverse as brain tumor, neurodegeneration and autoimmune diseases.

### Cell and stage-specific metabolic states

Metabolism and cell identity are closely coupled so that cells can carry out their function and meet the changing needs during the life cycle. Our transcriptomic data reveal the following key observations: a) all cell types show high levels of *ldh* expression in the embryo, b) they appear predominantly lipogenic in the embryo and c) they switch from being glycolytic in the embryo to being oxidative in the larva. While these features are shared by the three cell types across development, there are stark differences in terms of glucose uptake and subsequent break down. E16 hemocytes rely on *de novo* glucose and lipid biosynthesis and sustain a glycolytic state, while their larval counterpart depend more on the uptake of sugars and lipids and their subsequent breakdown to generate intermediates that run the TCA to sustain an oxidative state. E16 glia internalize sugars, generate lipids and metabolize them via glycolytic and oxidative routes, while L3 glia rely on gluconeogenesis and TAG biosynthesis and their breakdown to drive TCA and maintain an oxidative state. E16 neurons mediate sugar uptake and *de novo* lipogenesis and sustain a glycolytic state, whereas L3 neurons are more oxidative, which could be driven by *de novo* sugar synthesis and its breakdown to sustain TCA. L3 neurons also show reliance on TAG biosynthesis, which is perhaps critical for generating lipids essential for synaptic vesicle release. We speculate that in E16 neurons lipid synthesis and catabolism may function as a potential energy source while in E16 glia, lipid synthesis may be relevant for functions like secretory vesicular based intercellular communication.

In sum, embryonic cells are capable of intracellular synthesis of sugars and lipids and are also involved in internalizing extracellular metabolites. Later on, neurons and glia indulge in active uptake of metabolites and also continue to drive intracellular synthesis of lipids and sugars. Thus, both extracellular and intra-cellular resources drive the TCA in these tissues. By contrast, L3 hemocytes appear more reliant on extracellular nutrients to drive their internal lipid oxidation and TCA catabolic pathways. Such reliance on extracellular metabolites as the predominant source makes hemocytes competent sensors of systemic metabolites and any fluctuation in their levels.

The analyses of the metabolic pathways revealed an unexpected aspect of stage-specificity: the utilization of distinct paralogs in embryos and larvae. The fatty acid metabolic enzyme, *Fatty acid synthase 1 (FASN1, FBgn0027571)*, *CG3523,* is expressed in larvae, while its paralog *CG3524 (FASN2, FBgn0042627)*, was detected in embryos. The enzyme of lipid breakdown pathway, *CG17544 (FBgn0032775)* with predicted acyl-CoA oxidase activity, is expressed in larvae, while its paralog *CG9709, acyl-Coenzyme A oxidase at 57D distal (Acox57D-d, FBgn0034629)*, in embryos. Additionally, we identified genes that are not referred to as paralogs in Flybase, but share functional similarity, showing a similar trend: *CG6650 (FBgn0036402)*, predicted to have ADP-glucokinase activity, is expressed in larvae, while the glycolytic enzyme *CG8094*, *Hexokinase C* (*Hex-C, FBgn0001187*), in embryos. For the TCA enzymes, *CG7024 (FBgn0029722),* predicted to have pyruvate dehydrogenase activity, is expressed in larvae while *CG11876,* which is *Pyruvate dehydrogenase E1 beta subunit (Pdhb, FBgn0039635),* in embryos. The expression of *CG12233, Idh3a (FBgn0027291),* was detected in larvae, while *CG32026 (FBgn0052026),* predicted with isocitrate dehydrogenase activity, was noticed in embryos. Finally, the expression of *Ogdh (Fbgn0010352), CG11661,* was seen in embryos, while *CG5214 (FBgn0037891),* predicted to be part of the oxoglutarate dehydrogenase complex, in larvae. The enzymes with predicted acyl-CoA activity are tandemly duplicated, but the vast majority of the other genes are distantly located or reside on different chromosomes, calling for independent transcriptional regulation and/or stage-specific differences in their enzymatic activities. Future studies will establish the overall relevance of paralog switching on metabolic state transitions, but such temporal specificity provides an unexpected level of control that adds to the post transcriptional modifications of those enzymes.

The unbiased identification of common and specific transcriptional pathways allows a better understanding of the molecular processes associated to cell type identity. This genome-wide study has identified the transcriptional features of three types of related and unrelated cells, including their metabolic states and their transcriptional regulators. It has revealed novel cell-specific and stage-specific players including NimA and Rad50, respectively. It has highlighted time as a novel parameter defining a differentiated cell, paving the way to understand the role of high-order chromatin organization in the process. Altogether, this comparative high throughput analysis allows a better understanding of the mechanisms defining cell type identity, a central issue in biology and medical science.

## Materials and methods

### Fly strains and genetics

Flies were raised on standard medium at 25°C. Fly strains used are: *srp(hemo)Gal4/+; UAS-RFP/+* obtained upon crossing *srp(hemo)Gal4* (gift from K. Brückner) (Brückner et al., 2004) and *UAS-RFP* flies (RRID:BDSC_8547), *HmlΔ-RFP/+* (Makhijani et al., 2011), *srp(hemo)3xmcherry* (gift from D. Siekhaus, (Gyoergy et al., 2018), *repo-nRFP 43.1* on the 3rd chromosome (Laneve et al., 2013), *elav-nRFP 28.2* insertion on the 3rd chromosome *nSybGal4/+: UAS-RFP/+* obtained upon crossing *nSybGal4/+* (RRID:BDSC_51635) with *UAS-RFP/+*, *rad50Δ5.1* (Ciapponi et al., 2004) and *Oregon-R* strain was used as the control (referred to as *WT* in text). *NimAGal4* null mutant was generated by CRISPR-mediated mutagenesis and performed by WellGenetics Inc. using modified methods from (Kondo and Ueda, 2013). In brief, the upstream gRNA sequence ACTGCTCCTCCTGCTTGCAA[TGG] and downstream gRNA sequence GTGCCCTTCTAACATATACC[AGG] were cloned into U6 promote plasmid(s). Cassette *Gal4-3xP3-RFP*, which contains ribosome binding sequence (RBS), Gal4, SV40 polyA terminator and a floxed 3xP3-RFP, and two homology arms were cloned into pUC57-Kan as donor template for repair. *NimA*-targeting gRNAs and hs-Cas9 were supplied in DNA plasmids, together with donor plasmid for microinjection into embryos of control strain *w[1118]*. F1 flies carrying selection marker of 3xP3-RFP were further validated by genomic PCR and sequencing. CRISPR generates a 4,884-bp deletion removing the entire *NimA* CDS that is replaced by the cassette *Gal4-3xP3-RFP*. *NimAGal4; UAS-eGFP* animals were obtained upon crossing *NimAGal4 with UAS-eGFP*.

### FACS sorting of hemocytes, glia and neurons

Embryonic cells were isolated as follows. Staged egg laying was carried out to produce E16 embryos*. srp(hemo)Gal4/+; UAS-RFP/+*, *repo-nRFP*, *elav-nRFP* and *Oregon-R* strains were amplified, staged lays were done on yeast apple juice agar at 25°C. After a pre-lay period of 30 minutes, the agar plates and yeast were replaced with fresh plates and flies were left to lay for 3 hours at 25°C. Agar plates were removed and kept at 25°C, embryos were collected 11 hours and 40 minutes AEL, when they had reached stage 16. Embryos were then isolated from the medium and washed on a 100 µm mesh. The collected embryos were transferred into a cold solution of phosphate-buffered saline (PBS) in a Dounce homogenizer on ice. The embryos were dissociated using the large clearance pestle then the small clearance pestle, then filtered (70 µm filter).

L3 neurons, glia and hemocytes were purified from *elav-nRFP*, *repo-nRFP* and *HmlΔ-RFP/+* larvae, respectively. Staged lays of 3 hours were carried out at 25°C and wandering larvae were collected 108–117 hours AEL. L3 hemocytes were isolated as mentioned in (Cattenoz et al., 2020). Larvae were bled in cold PBS containing PTU (Sigma-Aldrich P7629) to prevent hemocyte melanization (Lerner and Fitzpatrick, 1950), and filtered (70 µm filter). For neurons and glia isolation, larval brains were dissected in cold PBS on ice and transferred to tubes containing 0.5 μg of collagenase IV (Gibco, Invitrogen) in 220 μL of PBS. Brains were incubated at 37°C on a Thermomixer with shaking at 500 rpm for 20 minutes. Then the brains were dissociated by pipetting up and down with 10-gauge needles and syringes and filtered through a 70 µm filter.

Cells were sorted using FACS Aria II (BD Biosciences) at 4°C in three independent biological replicates for each genotype. Live cells were first selected based on the forward scatter and side scatter and only single cells were sorted according to the RFP signal. *Oregon-R* cells were used as a negative control to set the gate and sort the RFP positive cells (**Sup. Figure 3-n**), which were collected directly in TRI reagent (MRC) for RNA extraction. Around 100 000 cells were sorted for each replicate. The purity of the sorted populations was assessed by carrying out a post-sort step. The FACS sorter was set up to produce cell pools displaying at least 80% of purity on the post-sort analysis.

### RNA extraction and sequencing

Sorted cells were homogenized in TRI reagent (MRC) for 5 minutes at room temperature (RT) to ensure complete dissociation of nucleoprotein complexes. 0.2 mL of chloroform was added to each sample followed by centrifugation at 12,000 g for 15 minutes at 4°C. The upper aqueous phase containing the RNA was collected and transferred to a new tube. 0.5 mL of 2-propanol were added, and the samples were incubated for 10 minutes at RT. The samples were then centrifuged at 12,000 g for 10 minutes to precipitate the RNA. The RNA pellet was then washed with 1 mL of 75% ethanol then precipitated again at 7,500 g for 5 minutes and air dried. 20 µL of RNase-free water was added to each sample before incubation at 55°C for 15 minutes. Single-end polyA+ RNA-Seq (mRNA-seq) libraries were prepared using the SMARTer (Takara) Low input RNA kit for Illumina sequencing. All samples were sequenced in 50-length Single-Read.

### Data analysis

The data analysis was done using the GalaxEast platform, the Galaxy instance of east of France (http://www.galaxeast.fr/). First, FastQC (Babraham Bioinformatics) was used on the raw FastQ files to generate summary statistics and assess the quality of the data. The raw files were then converted to FastQ Sanger using FastQ Groomer for downstream analysis. Data were then mapped to the *Drosophila melanogaster* August 2014 Dm6 (BDGF release 6 + ISO1 MT/Dm6) reference genome using TopHat (Trapnell et al., 2009). The number of reads per annotated gene were counted using Htseq-Count (Anders et al., 2015) and the comparison and normalization of the data between the different cell types was done using Deseq2 (Anders and Huber, 2010). To compare E16 stage to L3 using Deseq2, we considered all cell types deriving from E16 as replicates and compared to the cell deriving from L3 that were also considered replicates. Gene ontology studies were done using the Panther classification system version 16 (http://pantherdb.org/) (Mi et al., 2010; Thomas et al., 2003) for the identification of biological processes, Bonferroni correction was used for all analyses. Genes with a number of reads > 15 were considered significantly expressed. For the study of upregulated genes, only genes with fold change >1.5, adjusted p-value <0.05 and expression level >15 were taken into account. Genes with number of reads > 15 and adjusted p-value of fold change > 0.05 were considered non-differentially expressed. Amongst these, genes with number of reads > 15 and fold change < 1.5 were considered commonly expressed. For the study of scavenger receptors only genes with expression level > 500, adjusted p-value < 0.05 and fold change > 10 were considered as specific to either glia or hemocytes. The Venn diagrams were generated using Venny 2.1 (https://bioinfogp.cnb.csic.es/tools/venny/) (Oliveros, J.C. (2007-2015)) and data representation was done using Microsoft Excel. The heatmaps representing the log10 of the absolute expression levels in glia, neurons and hemocytes at E16 and L3 or representing the z-score were plotted in R (version 3.4.0) (R CoreTeam) using pheatmap (version 0.2) package (Kolde and Kolde, 2015).

### Quantitative PCR

For the comparison between the expression levels of scavenger receptors in hemocytes and glia, cells were isolated from *srp(hemo)-3xmCherry* and *repo-nRFP* wandering L3, respectively. FACS sorting and RNA extraction were as described above. The extracted RNA was then treated with DNase I recombinant RNase free (Roche) and the reverse transcription was done using the SuperScript IV (Invitrogen) with random primers. The cycle program used for the reverse transcription is 65°C for 10 minutes, 55°C for 20 minutes, 80°C for 10 minutes. The qPCR was done using SYBR Green I Master (Roche). Actin5C (Act5C) and Ribosomal protein 49 (RP49) were used to normalize the data. The primers are listed in **Sup. Table 9**. The p-values and statistical test used are indicated in the figure legends.

For the quantitative PCR on larval and embryonic tissues, whole larvae and whole embryos were crushed in cold PBS using a Dounce homogenizer, extracts were filtered first through a cell strainer of 100 μm and RNA extraction, reverse transcription and qPCR were done as described.

### Histone extraction and western blot

Histones from E16 and L3 *Oregon-R* were extracted as described in Abcam histone extraction protocol. Histone extracts were separated by 15% SDS-PAGE, transferred onto nitrocellulose membrane and probed with the primary antibodies mouse anti-H2A.v-P (1:5000) recognizing the *Drosophila* Histone 2A gamma variant, phosphorylated H2A.v-P (Developmental Studies Hybridoma Bank (DSHB AB_2618077, Lake et al., 2013) and rabbit anti-H3 (1:5000) (Abcam ab1791) for normalization. Signal was detected with Pierce ECL western blotting substrate (Thermo Fisher Scientific, Waltham, MA) using HRP-conjugated secondary antibodies (1:10000, Jackson).

### Immunolabeling of central nervous system

*NimAGal4; UAS-eGFP* embryos were dechorionated in bleach for 5 minutes and rinsed in water. They were then fixed in 50% heptane/50% paraformaldehyde (PFA) for 25 minutes on a shaker at RT and devitellinized in methanol/heptane for 1 minute. Embryos were then rinsed in methanol followed by PTX (0.3% Triton X-100 in PBS 1x) and incubated in blocking reagent (Roche) for 1 hour at RT. They were incubated with primary antibodies mouse anti-Repo 1:20 (DSHB #8D12), chicken anti-GFP 1:500 (Abcam ab13970) and rabbit anti-Nazgul 1:100 (gift From B. Altenhein) in blocking reagent overnight (O/N) at 4°C, then washed in PTX 3 × 10 minutes at RT and incubated with secondary antibodies 1:400 (Cy3 anti-mouse and FITC anti-chicken (Jackson)) for 1 hour at RT and washed 3x 10 minutes at PTX. The embryos were mounted in Vectashield (Vector #H-1000) and analyzed by confocal microscopy (Leica spinning disk). For L3 larvae and adults, CNSs from *NimAGal4; UAS-eGFP* were dissected in cold PBS 1x and transferred into wells containing 4% PFA and fixed for 2 hours at RT. They were then washed in PTX for 1 hour and incubated in blocking reagent for 1 hour at RT. CNSs were then incubated with primary antibodies mouse anti-Repo, chicken anti-GFP and rabbit anti-Nazgul O/N at 4°C, then washed in PTX 3x 10 minutes and incubated with secondary antibodies (Cy5 anti-mouse, Cy3 anti-rabbit (Jackson)) for 1 hour at RT. After washing in PTX, CNSs were mounted in Vectashield imaged using Leica spinning disk microscope and analyzed using Fiji (National Institute of Mental Health, Bethesda, Maryland, USA, (Schindelin et al., 2012).

### Immunolabeling of polytene chromosomes

Salivary glands were dissected from L3 larvae in NaCl 0.7% / NP-40 0.1% and fixed for 2 minutes in 3.7% formaldehyde/ 0.7% NaCl/ 1% Triton-X. Before squashing, the glands were transferred in 2% formaldehyde/ 45% acetic acid and incubated for 2 minutes on a siliconized coverslip. During incubation, salivary glands were fragmented. A poly-L-lysine slide was inverted on top of the coverslip and salivary glands were squashed. The chromosome spreads were examined by phase microscopy and frozen in liquid nitrogen. The coverslip was flicked off with a razor blade and slides were incubated in cold PBS 2× for 15 min. The slides were transferred to a blocking solution (3% BSA, 2% NP-40, 0.2% Tween-20, 10% non-fat dry milk in PBS) and blocked for 1 hour at RT. After blocking, slides were incubated with primary antibody in blocking solution overnight at 4°C in a humid chamber. Slides were then rinsed in PBS and washed 2x 15 minutes in PBS1x /0.2% Tween/0.1% NP-40. After washing, slides were rinsed in PBS and incubated with secondary antibody in blocking solution 2 hours at RT in a humid chamber. Slides were then rinsed in PBS and washed 2x 15 minutes in PBS1x /0.2% Tween/0.1% NP-40. Finally, slides were drained and mounted with Vectashield medium H-1200 with DAPI to stain DNA. The primary antibodies used for immunolabeling were: rabbit anti-H3K4me3 1:400 (Abcam ab8580); mouse anti-H3K9me2 1:20 (Abcam ab1220); rabbit anti-H3K9me3 1:100 (Abcam ab8898); rabbit anti-H3K27me3 1:400 (Cell Signaling C36B11); mouse anti-H2A.v-P 1:20 (DSHB AB_2618077). The secondary antibodies were: FITC-conjugated goat anti-rabbit 1:50 (Jackson) and Rhodamine conjugated goat anti-mouse 1:50 (Jackson). Polytene chromosomes preparations were analyzed on a fluorescence microscope (Zeiss Apotome) and images acquisition performed using the Zen Pro software (Zeiss). Polytene chromosome fluorescence intensity and bands plot profile analysis were performed using Fiji software (National Institute of Mental Health, Bethesda, Maryland, USA).

### *In situ* hybridization

We used a RNAscope® probe and reagent kit. Larval CNSs from *Oregon R* were dissected in cold PBS 1x and fixed in 4% PFA O/N at 4°C then washed 3x 10 minutes in PBST (PBS 1x/ 0.1% Tween-20) followed by one wash in PBS 1x. Samples were then incubated in RNAscope reagents as in protease III at 40°C for 10 minutes and washed in 3x 2 minutes in PBS 1x. The probe targeting *NimA* was then added in addition to the positive control probe and samples were incubated O/N at 40°C. The amplifiers and detection solution in the RNAscope reagent kit were added sequentially for hybridization signal amplification for the indicated time. CNSs were then mounted in Vectashield, imaged using Leica spinning disk microscope and analyzed using Fiji.

### Metabolic pathway analysis

All data points considered for our analysis has a read count of more than ten in at least one of the samples from the tissue under consideration. Differential expression analysis between the L3 and E16 developmental stages was done for each tissue using DESeq2. Genes associated with metabolic pathways considered in this study, were retrieved from the KEGG database (Kanehisa and Goto, 2000, http://www.genome.jp/). The heatmaps representing the z-score of the metabolic genes in glia, neurons and hemocytes at E16 and L3 were plotted using R (version 3.4.0) (R CoreTeam). For genes with paralogs, the paralogs show log2FC(L3/E16) greater or lesser than zero, the paralog with the highest fold-change has been plotted. For genes whose paralogs have log2FC(L3/E16) that is either positive or negative, the paralogs with the highest and lowest fold-change have been plotted. The details of all the genes (including all the paralogs) as well as genes included in the figures are present in **Sup. Table 7.**

## Acknowledgments

We thank the Imaging Center of the IGBMC for technical assistance. The sequencing was performed by the GenomEast platform, a member of the “France Génomique” consortium (ANR-10-INBS-0009). We thank K. Bruckner, D. Siekhaus and B. Altenhein for providing fly stocks and antibodies. In addition, stocks obtained from the Bloomington *Drosophila* Stock Center (NIH P40OD018537) and antibodies obtained from the Developmental Studies Hybridoma Bank created by the NICHD of the NIH and maintained at The University of Iowa (Department of Biology, Iowa City, IA 52242) were used in this study. This work was supported by INSERM, CNRS, UDS, Ligue Régionale contre le Cancer, Hôpital de Strasbourg, ARC, CEFIPRA, USIAS, FRM and ANR grants, and by the CNRS/University LIA Calim. R. Sakr was supported by the French state fund through a doctoral contract from the University of Strasbourg and the Fondation pour la Recherche Médicale (FDT2020010107630). The IGBMC was also supported by a French state fund through the ANR labex. L. Ciapponi’s lab Sapienza University grant RM120172B7D32C04. T. Mukherjee’s lab is supported by the DST-Core Reasearch Grant, DBT-IYBA 2017, CEFIPRA and USIAS fellowship. NH is a Graduate Student at inStem and supported by Council for Scientific and Industrial Research fellowship (CSIR).

## Author contribution

Conceptualization, RS and AG; Methodology, RS, AG, PBC, RS, AG, PBC, AP, MM, LC, TM and NH. Investigation, RS, AG, PBC, AP, MM, LC, TM and NH. Writing—Original Draft, RS and AG, Writing—Review and Editing, RS, AG and PBC; Funding Acquisition, AG, LC and TM; Resources, AG; Supervision, AG and PBC.

## Supplementary figure legends

**Sup. Figure 1:**
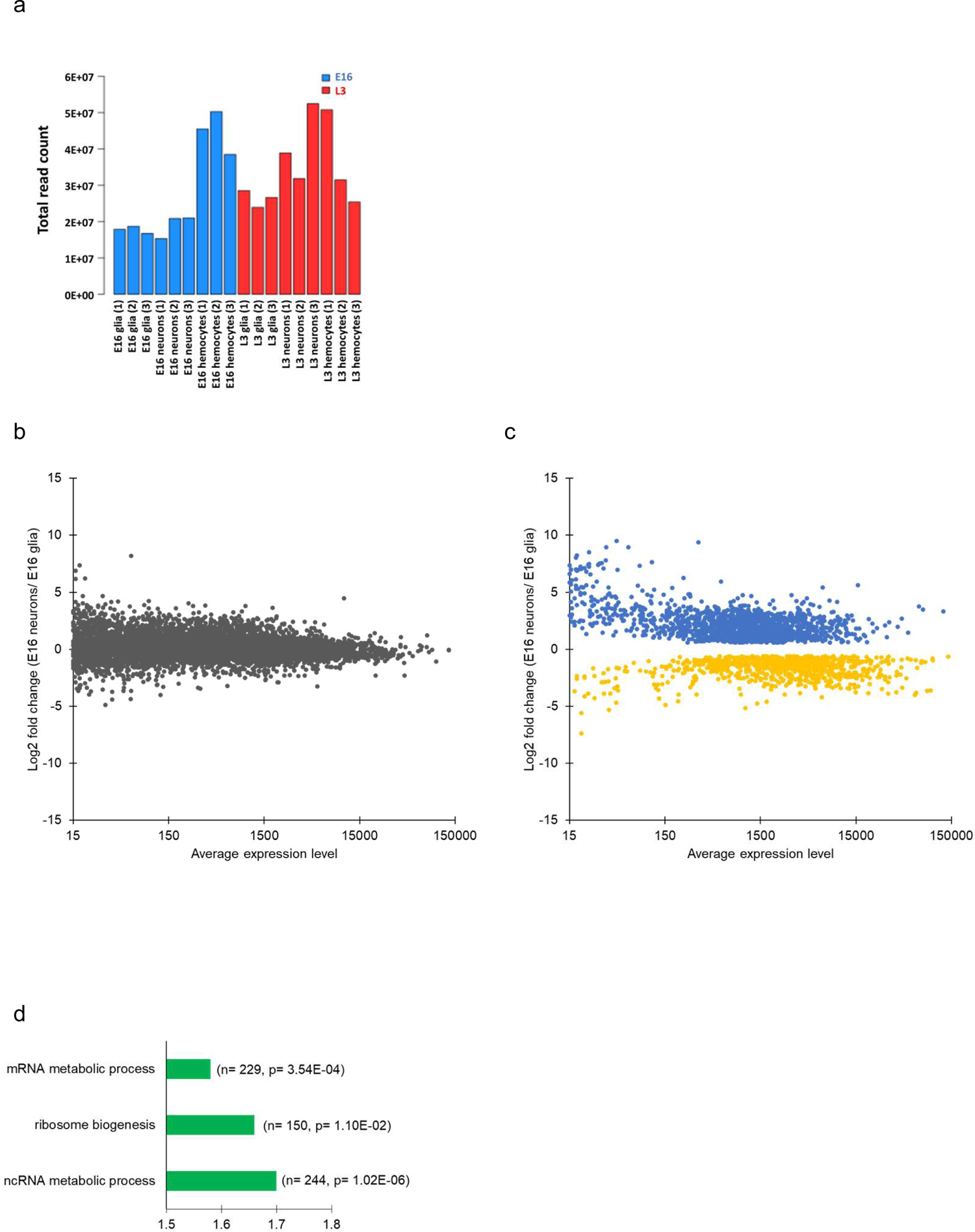
Read counts and commonalities between embryonic neurons and glia. (a) Comparison of the total read count between the different samples. The y-axis represents the total read count. The x axis-represents the different samples: E16 glia, neurons and hemocytes (blue) and L3 glia, neurons and hemocytes (red). (b,c) Dot plots comparing the expression levels of non-differentially genes versus differentially expressed genes. y-axis reports the log2 fold change E16 neurons/E16 glia. Non-differentially expressed genes are shown in gray (b). The x-axis reports the average gene expression levels. Genes significantly upregulated in E16 neurons are shown in blue, genes significantly upregulated in E16 glia are shown in yellow (c). (d) Gene Ontology (GO) term enrichment analysis on genes commonly expressed between E16 glia and neurons. The fold enrichment for a subset of representative GO terms is displayed on the x-axis, the number of genes and the P-value of the GO term enrichment are indicated in brackets. The complete results of the GO term analysis are shown in **Sup. Table 2-a.**

**Sup. Figure 2:**
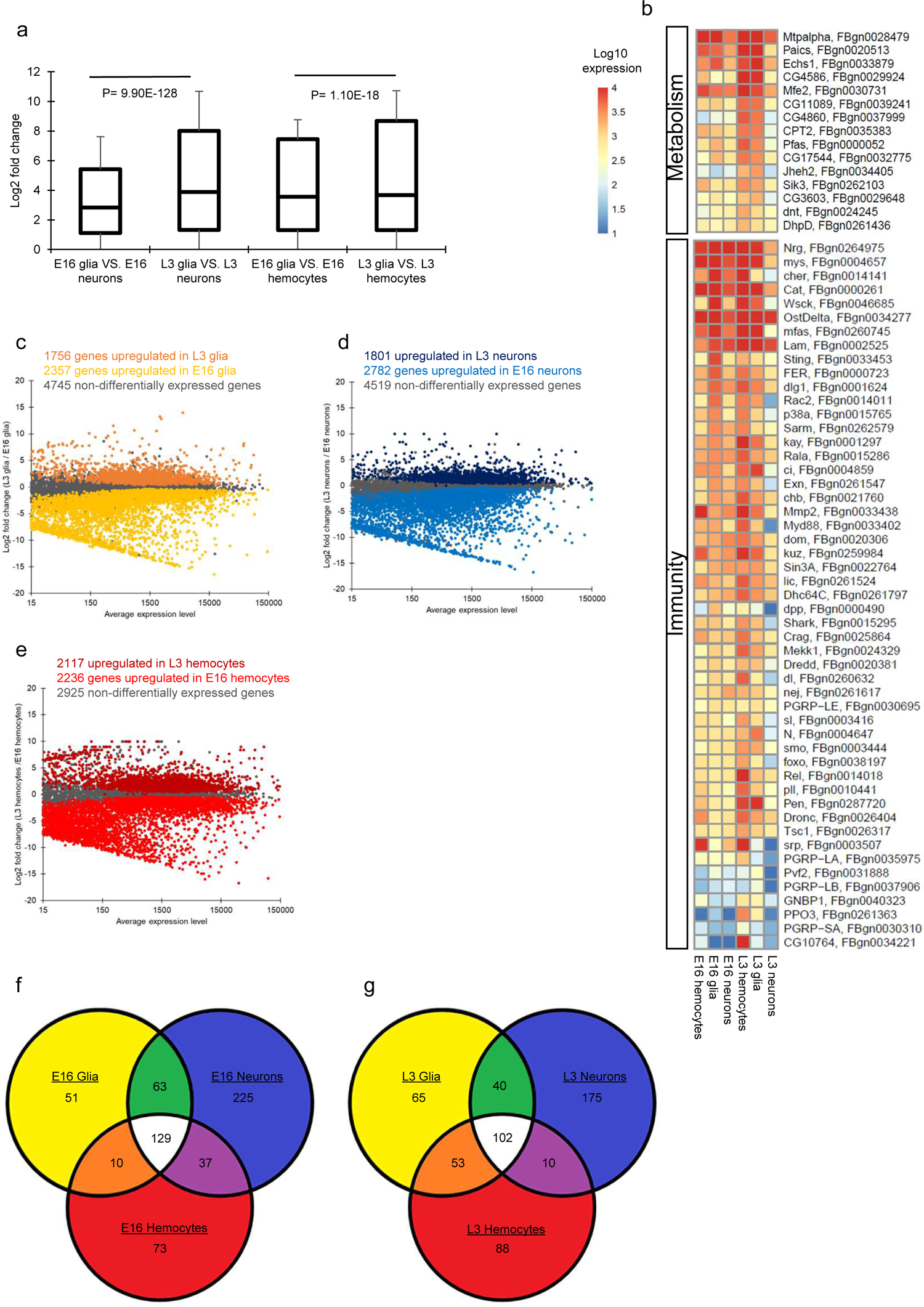
Cell type differences between E16 and L3. (a) Box plot showing the log2 fold change (y-axis) between the different samples at E16 and L3 (x-axis). P = p-value calculated using Student’s t-test and indicated on graphs, error bars represent the standard deviation. (b) Heatmap showing the Log10 expression levels (absolute levels) of genes upregulated in L3 hemocytes and L3 glia but not in neurons and involved in metabolism. Scale shows low expression (blue) to high expression (red). Log10 expression values are shown in **Sup. Table 4-b,c**. (c-e) Transcriptome comparison of the cell types at E16 vs. L3. The x-axis reports the average gene expression levels, the y-axis, the log2 fold change of L3 glia / E16 glia (c), L3 neurons / E16 neurons (d) and L3 hemocytes / E16 hemocytes (e). Genes significantly upregulated in L3 glia, L3 neurons and L3 hemocytes are shown in orange, dark blue and dark red respectively. Genes significantly upregulated in E16 glia, E16 neurons and E16 hemocytes are shown in yellow, light blue and light red, respectively. Non-differentially expressed genes are shown in gray. (f,g) Venn diagrams showing the distribution of transcription factors between E16 neurons (blue), glia (yellow) and hemocytes (red) (f), and L3 neurons (blue), glia (yellow) and hemocytes (red) (g). Transcription factors common to all three cell types are shown in white. The number of transcription factors in each category is indicated in the diagram.

**Sup. Figure 3:**
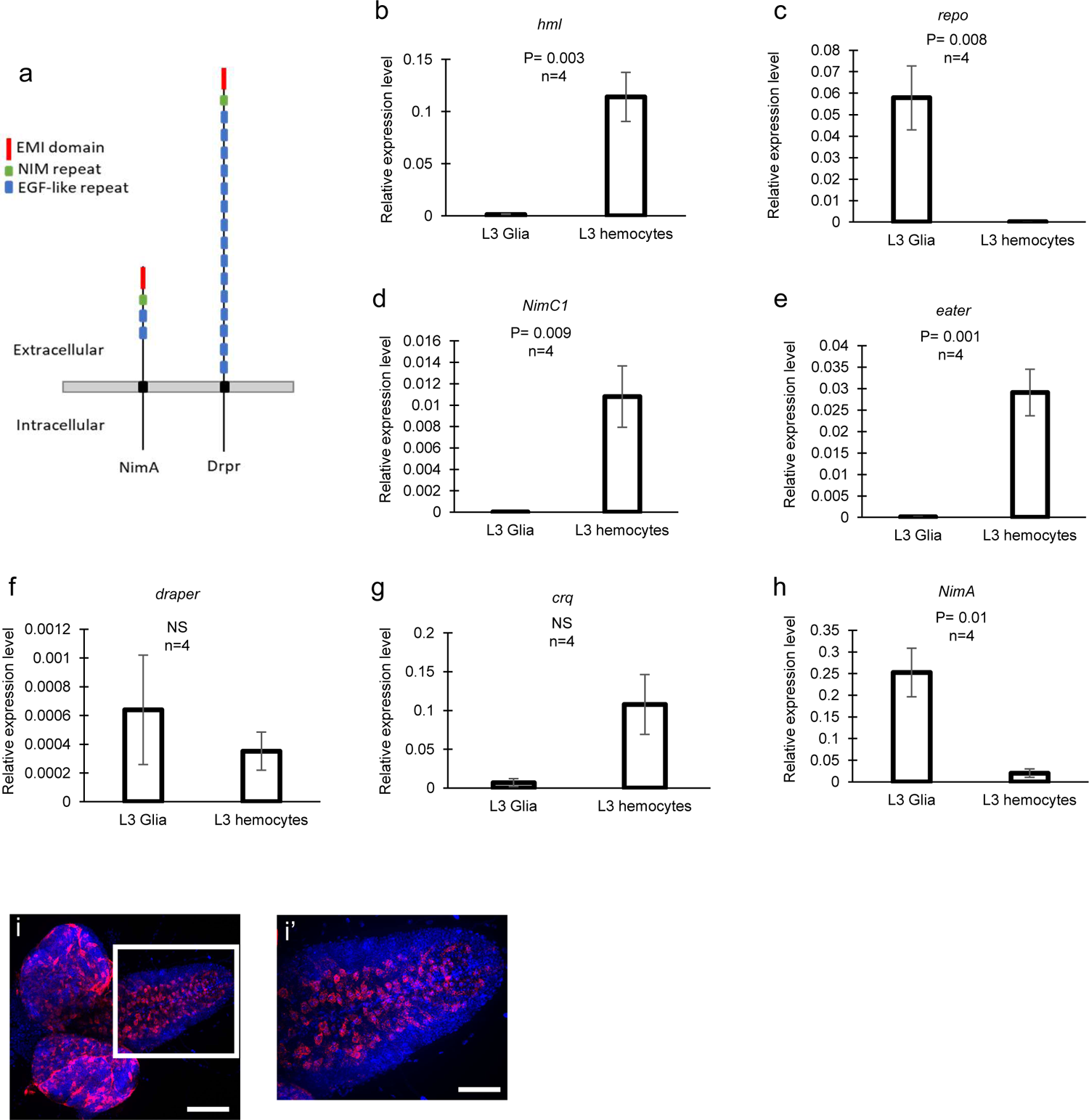
Commonalities and specificities between hemocytes and glia. (a) Graphical representation of NimA and Drpr. (b-h) Quantitative reverse transcription PCR (RT-qPCR) assays on sorted L3 glia and hemocytes. To test the purity of the sorted populations, the expression of *Hml* (b) and *repo* (c) were assessed in hemocytes and glia, respectively. The levels of *NimC1* (d), *eater* (e), *drpr* (f), *crq* (g) and *NimA* (h) were quantified and relative expression levels were calculated based on the expression of *Act5C* and *RP49*. List of primers shown in **Sup. Table 9**, n = number of biological replicates, P = p-value calculated using Student’s t-test and indicated on graphs, error bars represent the standard deviation. NS = non-significant. (i) *In situ* hybridization using an anti-*NimA* RNAscope probe on L3 CNS, full stack projection. Region in white square in (i) is shown magnified in (i’). *NimA* is shown in red, DAPI in blue. (i) Scale bar = 100 μm, (i’) Scale bar = 50 μm.

**Sup. Figure 4:**
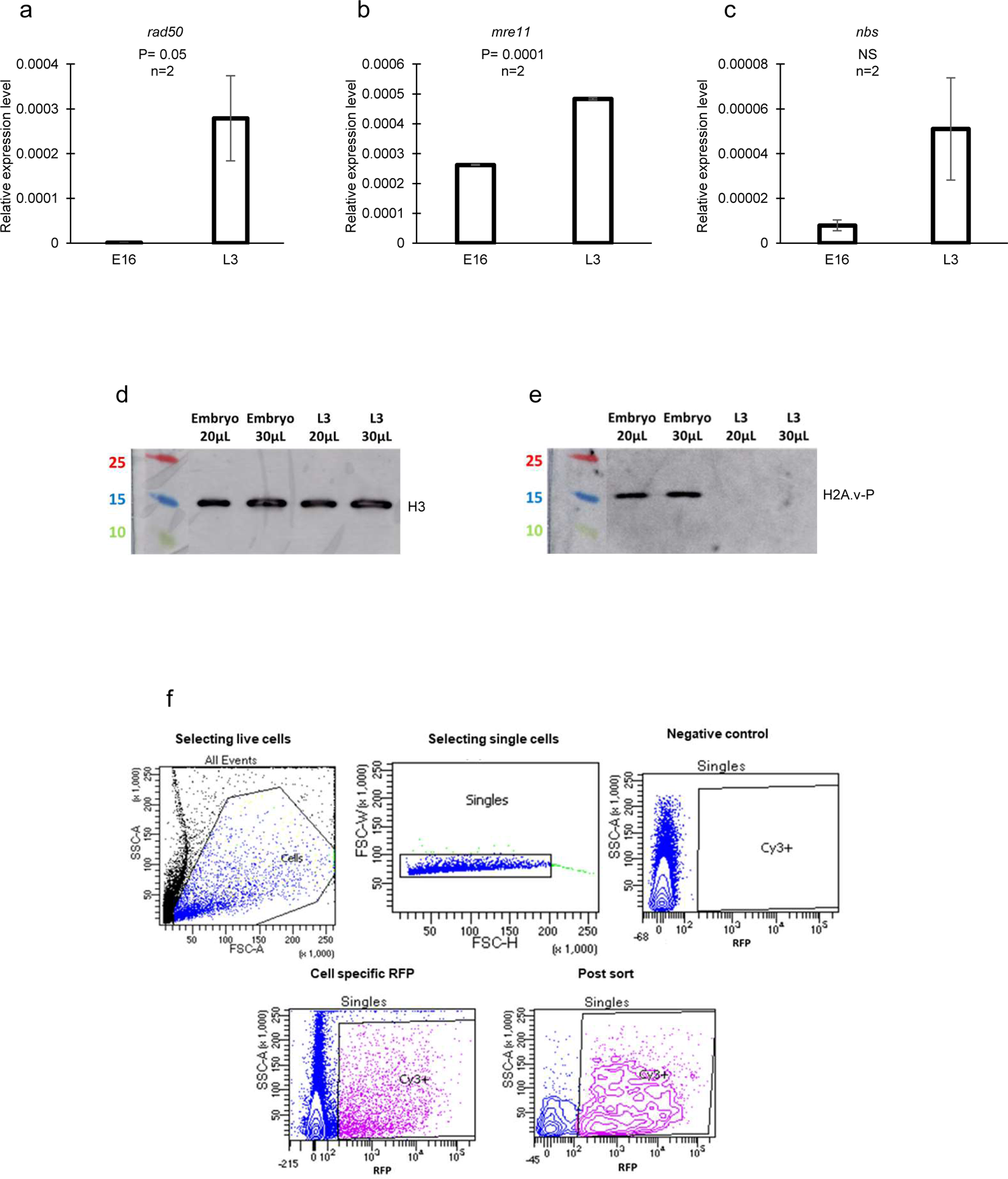
Specificities of larval cells. (a-c) RT-qPCR assays on cells from E16 and L3 *WT* animals. The levels of *rad50* (a), *mre11* (b) and *nbs* (c) transcripts were quantified and the relative expression levels were calculated based on the expression of *Act5C* and *RP49.* List of primers shown in **Sup. Table 9**, n = number of biological replicates, p-value calculated using Student’s t-test and indicated on graphs, error bars represent the standard deviation. (d,e) Western blot quantification of γH2AV. Histones extracted from cells from E16 and L3 *WT* animals were blotted on nitrocellulose membranes and probed with anti-H2A.v-P antibody (e). An antibody that recognizes all histones H3, anti-H3, was used to confirm equal loading of lysates (d). The ladder on the left shows protein size in KDa. (f) Cells were first selected based on the forward scatter (FSC-A) and the side scatter (SSC-A) to eliminate dead cells and debris. Doublets were removed and only single cells were taken into account. The gate for sorting Cy3/RFP positive cells was based on the negative control. Only Cy3/RFP positive were sorted. A post sort step was performed at the end to test the purity of the sorted population.

**Sup. Figure 5:**
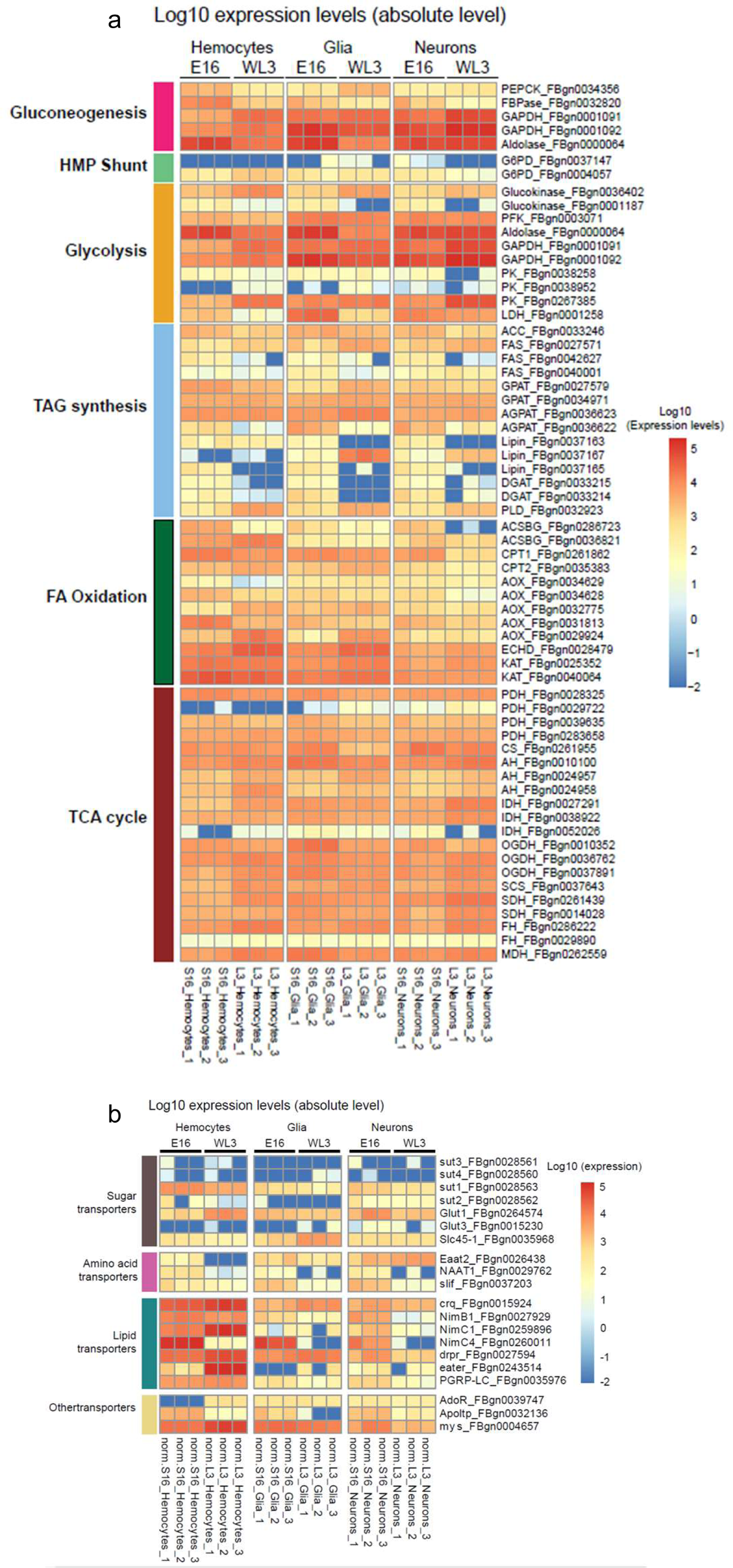
Comparison of the metabolic pathways expressed in the three cell types at embryonic and larval stages. (a,b) Heatmaps showing the Log10 expression levels (absolute levels) of genes involved in the main metabolic processes (a) and of the main transporters (b) in hemocytes, glia and neurons of E16 and L3. Scale shows low expression (blue) to high expression (red). Expression values are shown in **Sup. Table 7**.

## List of supplementary tables

**Sup. Table 1:** List of transcription factors and their expression levels in E16 and L3 glia, neurons and hemocytes.

**Sup. Table 2:** Gene Ontology terms found in commonly expressed genes or genes upregulated in E16 glia, E16 neurons or E16 hemocytes.

**Sup. Table 3:** Gene Ontology terms found in genes upregulated in L3 glia, L3 neurons or L3 hemocytes.

**Sup. Table 4:** Gene Ontology terms and list of genes upregulated in L3 glia and hemocytes compared to neurons.

**Sup. Table 5:** List of scavenger receptors expressed in glia and hemocytes.

**Sup. Table 6:** Gene Ontology terms and lists of genes upregulated in E16 cells or L3 cells.

**Sup. Table 7:** Lists of genes involved in different metabolism pathways.

**Sup. Table 8:** Gene Ontology terms found in genes upregulated in E16 hemocytes or E16 neurons.

**Sup. Table 9:** List of primers used in this study.

## References

1. Abbott, N.J. (2005). Dynamics of CNS barriers: evolution, differentiation, and modulation. Cell. Mol. Neurobiol. 25, 5–23.

2. Anders, S., and Huber, W. (2010). Differential expression analysis for sequence count data. Genome Biol. 11.

3. Anders, S., Pyl, P.T., and Huber, W. (2015). HTSeq--a Python framework to work with high-throughput sequencing data. Bioinformatics 31, 166–169.

4. Aranda, S., Mas, G., and Di Croce, L. (2015). Regulation of gene transcription by Polycomb proteins. Sci. Adv. 1, e1500737.

5. Awasaki, T., and Ito, K. (2004). Engulfing action of glial cells is required for programmed axon pruning during Drosophila metamorphosis. Curr. Biol. 14, 668–677.

6. Barres, B.A. (2008). The mystery and magic of glia: a perspective on their roles in health and disease. Neuron 60, 430–440.

7. Basenko, E.Y., Sasaki, T., Ji, L., Prybol, C.J., Burckhardt, R.M., Schmitz, R.J., and Lewis, Z.A. (2015). Genome-wide redistribution of H3K27me3 is linked to genotoxic stress and defective growth. Proc. Natl. Acad. Sci. U. S. A. 112, E6339–48.

8. Bittern, J., Pogodalla, N., Ohm, H., Brüser, L., Kottmeier, R., Schirmeier, S., and Klämbt, C. (2021). Neuron-glia interaction in the Drosophila nervous system. Dev. Neurobiol. 81, 438–452.

9. Bosso, G., Cipressa, F., Moroni, M.L., Pennisi, R., Albanesi, J., Brandi, V., Cugusi, S., Renda, F., Ciapponi, L., Polticelli, F., et al. (2019). NBS1 interacts with HP1 to ensure genome integrity. Cell Death Dis. 10, 951.

10. Brink, D.L., Gilbert, M., Xie, X., Petley-Ragan, L., and Auld, V.J. (2012). Glial processes at the Drosophila larval neuromuscular junction match synaptic growth. PLoS One 7, e37876.

11. Brown, N.H. (2011). Extracellular matrix in development: insights from mechanisms conserved between invertebrates and vertebrates. Cold Spring Harb. Perspect. Biol. 3.

12. Brückner, K., Kockel, L., Duchek, P., Luque, C.M., Rørth, P., and Perrimon, N. (2004). The PDGF/VEGF receptor controls blood cell survival in Drosophila. Dev. Cell 7, 73–84.

13. Callebaut, I., Mignotte, V., Souchet, M., and Mornon, J.P. (2003). EMI domains are widespread and reveal the probable orthologs of the Caenorhabditis elegans CED-1 protein. Biochem. Biophys. Res. Commun. 300, 619–623.

14. Cattenoz, P.B., Sakr, R., Pavlidaki, A., Delaporte, C., Riba, A., Molina, N., Hariharan, N., Mukherjee, T., and Giangrande, A. (2020). Temporal specificity and heterogeneity of Drosophila immune cells. EMBO J. e104486.

15. Ciapponi, L., Cenci, G., Ducau, J., Flores, C., Johnson-Schlitz, D., Gorski, M.M., Engels, W.R., and Gatti, M. (2004). The Drosophila Mre11/Rad50 complex is required to prevent both telomeric fusion and chromosome breakage. Curr. Biol. 14, 1360–1366.

16. Ciapponi, L., Cenci, G., and Gatti, M. (2006). The Drosophila Nbs protein functions in multiple pathways for the maintenance of genome stability. Genetics 173, 1447–1454.

17. Conway, E., Healy, E., and Bracken, A.P. (2015). PRC2 mediated H3K27 methylations in cellular identity and cancer. Curr. Opin. Cell Biol. 37, 42–48.

18. Cornman, R.S. (2009). Molecular evolution of Drosophila cuticular protein genes. PLoS One 4, e8345.

19. Daines, B., Wang, H., Wang, L., Li, Y., Han, Y., Emmert, D., Gelbart, W., Wang, X., Li, W., Gibbs, R., et al. (2011). The Drosophila melanogaster transcriptome by paired-end RNA sequencing. Genome Res. 21, 315–324.

20. Davie, K., Janssens, J., Koldere, D., De Waegeneer, M., Pech, U., Kreft, Ł., Aibar, S., Makhzami, S., Christiaens, V., Bravo González-Blas, C., et al. (2018). A Single-Cell Transcriptome Atlas of the Aging Drosophila Brain. Cell 174, 982–998.e20.

21. Deliu, L.P., Ghosh, A., and Grewal, S.S. (2017). Investigation of protein synthesis in Drosophila larvae using puromycin labelling. Biol. Open 6, 1229–1234.

22. DePaula-Silva, A.B., Gorbea, C., Doty, D.J., Libbey, J.E., Sanchez, J.M.S., Hanak, T.J., Cazalla, D., and Fujinami, R.S. (2019). Differential transcriptional profiles identify microglial- and macrophage-specific gene markers expressed during virus-induced neuroinflammation. J. Neuroinflammation 16, 1–20.

23. Ebens, A.J., Garren, H., Cheyette, B.N., and Zipursky, S.L. (1993). The Drosophila anachronism locus: a glycoprotein secreted by glia inhibits neuroblast proliferation. Cell 74, 15–27.

24. Forey, R., Barthe, A., Tittel-Elmer, M., Wery, M., Barrault, M.-B., Ducrot, C., Seeber, A., Krietenstein, N., Szachnowski, U., Skrzypczak, M., et al. (2021). A Role for the Mre11-Rad50-Xrs2 Complex in Gene Expression and Chromosome Organization. Mol. Cell 81, 183–197.e6.

25. Franc, N.C., Dimarcq, J.L., Lagueux, M., Hoffmann, J., and Ezekowitz, R.A. (1996). Croquemort, a novel Drosophila hemocyte/macrophage receptor that recognizes apoptotic cells. Immunity 4, 431–443.

26. Franc, N.C., Heitzler, P., Ezekowitz, R.A., and White, K. (1999). Requirement for croquemort in phagocytosis of apoptotic cells in Drosophila. Science 284, 1991–1994.

27. Garces, A., and Thor, S. (2006). Specification of Drosophila aCC motoneuron identity by a genetic cascade involving even-skipped, grain and zfh1. Development 133, 1445–1455.

28. Gatei, M., Kijas, A.W., Biard, D., Dörk, T., and Lavin, M.F. (2014). RAD50 phosphorylation promotes ATR downstream signaling and DNA restart following replication stress. Hum. Mol. Genet. 23, 4232– 4248.

29. Gyoergy, A., Roblek, M., Ratheesh, A., Valoskova, K., Belyaeva, V., Wachner, S., Matsubayashi, Y., Sánchez-Sánchez, B.J., Stramer, B., and Siekhaus, D.E. (2018). Tools Allowing Independent Visualization and Genetic Manipulation of Drosophila melanogaster Macrophages and Surrounding Tissues. G3 (Bethesda). 8, 845–857.

30. Hickman, S.E., Kingery, N.D., Ohsumi, T.K., Borowsky, M.L., Wang, L.C., Means, T.K., and El Khoury, J. (2013). The microglial sensome revealed by direct RNA sequencing. Nat. Neurosci. 2013 1612 16, 1896– 1905.

31. Hildebrandt, A., Pflanz, R., Behr, M., Tarp, T., Riedel, D., and Schuh, R. (2015). Bark beetle controls epithelial morphogenesis by septate junction maturation in Drosophila. Dev. Biol. 400, 237–247.

32. James, T.C., and Elgin, S.C. (1986). Identification of a nonhistone chromosomal protein associated with heterochromatin in Drosophila melanogaster and its gene. Mol. Cell. Biol. 6, 3862–3872.

33. James, T.C., Eissenberg, J.C., Craig, C., Dietrich, V., Hobson, A., and Elgin, S.C. (1989). Distribution patterns of HP1, a heterochromatin-associated nonhistone chromosomal protein of Drosophila. Eur. J. Cell Biol. 50, 170–180.

34. Kanehisa, M., and Goto, S. (2000). KEGG: kyoto encyclopedia of genes and genomes. Nucleic Acids Res. 28, 27–30.

35. Karouzou, M. V, Spyropoulos, Y., Iconomidou, V.A., Cornman, R.S., Hamodrakas, S.J., and Willis, J.H. (2007). Drosophila cuticular proteins with the R&R Consensus: annotation and classification with a new tool for discriminating RR-1 and RR-2 sequences. Insect Biochem. Mol. Biol. 37, 754–760.

36. Kocks, C., Cho, J.H., Nehme, N., Ulvila, J., Pearson, A.M., Meister, M., Strom, C., Conto, S.L., Hetru, C., Stuart, L.M., et al. (2005). Eater, a transmembrane protein mediating phagocytosis of bacterial pathogens in Drosophila. Cell 123, 335–346.

37. Kondo, S., and Ueda, R. (2013). Highly improved gene targeting by germline-specific Cas9 expression in Drosophila. Genetics 195, 715–721.

38. Kurant, E., Axelrod, S., Leaman, D., and Gaul, U. (2008). Six-microns-under acts upstream of Draper in the glial phagocytosis of apoptotic neurons. Cell 133, 498–509.

39. Kurucz, E., Márkus, R., Zsámboki, J., Folkl-Medzihradszky, K., Darula, Z., Vilmos, P., Udvardy, A., Krausz, I., Lukacsovich, T., Gateff, E., et al. (2007). Nimrod, a putative phagocytosis receptor with EGF repeats in Drosophila plasmatocytes. Curr. Biol. 17, 649–654.

40. Lake, C.M., Holsclaw, J.K., Bellendir, S.P., Sekelsky, J., and Hawley, R.S. (2013). The development of a monoclonal antibody recognizing the Drosophila melanogaster phosphorylated histone H2A variant (γ-H2AV). G3 (Bethesda). 3, 1539–1543.

41. Landgraf, M., and Thor, S. (2006). Development of Drosophila motoneurons: specification and morphology. Semin. Cell Dev. Biol. 17, 3–11.

42. Laneve, P., Delaporte, C., Trebuchet, G., Komonyi, O., Flici, H., Popkova, A., D’Agostino, G., Taglini, F., Kerekes, I., and Giangrande, A. (2013). The Gcm/Glide molecular and cellular pathway: new actors and new lineages. Dev. Biol. 375, 65–78.

43. Lebestky, T., Chang, T., Hartenstein, V., and Banerjee, U. (2000). Specification of Drosophila hematopoietic lineage by conserved transcription factors. Science 288, 146–149.

44. Lemaitre, B., Nicolas, E., Michaut, L., Reichhart, J.M., and Hoffmann, J.A. (1996). The dorsoventral regulatory gene cassette spätzle/Toll/cactus controls the potent antifungal response in Drosophila adults. Cell 86, 973–983.

45. Lerner, A.B., and Fitzpatrick, T.B. (1950). Biochemistry of melanin formation. Physiol. Rev. 30, 91–126.

46. Li, Q., and Barres, B.A. (2017). Microglia and macrophages in brain homeostasis and disease. Nat. Rev. Immunol. 2017 184 18, 225–242.

47. Limmer, S., Weiler, A., Volkenhoff, A., Babatz, F., and Klambt, C. (2014). The Drosophila blood-brain barrier: development and function of a glial endothelium. Front. Neurosci. 8, 365.

48. MacDonald, J.M., Beach, M.G., Porpiglia, E., Sheehan, A.E., Watts, R.J., and Freeman, M.R. (2006). The Drosophila cell corpse engulfment receptor Draper mediates glial clearance of severed axons. Neuron 50, 869–881.

49. Makhijani, K., Alexander, B., Tanaka, T., Rulifson, E., and Brückner, K. (2011). The peripheral nervous system supports blood cell homing and survival in the Drosophila larva. Development 138, 5379–5391.

50. Manaka, J., Kuraishi, T., Shiratsuchi, A., Nakai, Y., Higashida, H., Henson, P., and Nakanishi, Y. (2004). Draper-mediated and phosphatidylserine-independent phagocytosis of apoptotic cells by Drosophila hemocytes/macrophages. J. Biol. Chem. 279, 48466–48476.

51. Mangahas, P.M., and Zhou, Z. (2005). Clearance of apoptotic cells in Caenorhabditis elegans. Semin. Cell Dev. Biol. 16, 295–306.

52. Martinek, N., Shahab, J., Saathoff, M., and Ringuette, M. (2008). Haemocyte-derived SPARC is required for collagen-IV-dependent stability of basal laminae in Drosophila embryos. J. Cell Sci. 121, 1671–1680.

53. Melcarne, C., Lemaitre, B., and Kurant, E. (2019). Phagocytosis in Drosophila: From molecules and cellular machinery to physiology. Insect Biochem. Mol. Biol. 109, 1–12.

54. Mi, H., Dong, Q., Muruganujan, A., Gaudet, P., Lewis, S., and Thomas, P.D. (2010). PANTHER version 7: improved phylogenetic trees, orthologs and collaboration with the Gene Ontology Consortium. Nucleic Acids Res. 38.

55. Naba, A., Clauser, K.R., Ding, H., Whittaker, C.A., Carr, S.A., and Hynes, R.O. (2016). The extracellular matrix: Tools and insights for the “omics” era. Matrix Biol. 49, 10–24.

56. Neumann, H., Kotter, M.R., and Franklin, R.J.M. (2009). Debris clearance by microglia: an essential link between degeneration and regeneration. Brain 132, 288–295.

57. O’Hagan, H.M., Mohammad, H.P., and Baylin, S.B. (2008). Double strand breaks can initiate gene silencing and SIRT1-dependent onset of DNA methylation in an exogenous promoter CpG island. PLoS Genet. 4, e1000155.

58. O’Hagan, H.M., Wang, W., Sen, S., Destefano Shields, C., Lee, S.S., Zhang, Y.W., Clements, E.G., Cai, Y., Van Neste, L., Easwaran, H., et al. (2011). Oxidative damage targets complexes containing DNA methyltransferases, SIRT1, and polycomb members to promoter CpG Islands. Cancer Cell 20, 606–619.

59. Ou, J., He, Y., Xiao, X., Yu, T.-M., Chen, C., Gao, Z., and Ho, M.S. (2014). Glial cells in neuronal development: recent advances and insights from Drosophila melanogaster. Neurosci. Bull. 30, 584–594.

60. Parker, R.J., and Auld, V.J. (2006). Roles of glia in the Drosophila nervous system. Semin. Cell Dev. Biol. 17, 66–77.

61. Pearson, A., Lux, A., and Krieger, M. (1995). Expression cloning of dSR-CI, a class C macrophage-specific scavenger receptor from Drosophila melanogaster. Proc. Natl. Acad. Sci. U. S. A. 92, 4056–4060.

62. Peco, E., Davla, S., Camp, D., Stacey, S.M., Landgraf, M., and van Meyel, D.J. (2016). Drosophila astrocytes cover specific territories of the CNS neuropil and are instructed to differentiate by Prospero, a key effector of Notch. Development 143, 1170–1181.

63. Pereanu, W., Shy, D., and Hartenstein, V. (2005). Morphogenesis and proliferation of the larval brain glia in Drosophila. Dev. Biol. 283, 191–203.

64. Peters, A.H.F.M., Kubicek, S., Mechtler, K., O’Sullivan, R.J., Derijck, A.A.H.A., Perez-Burgos, L., Kohlmaier, A., Opravil, S., Tachibana, M., Shinkai, Y., et al. (2003). Partitioning and plasticity of repressive histone methylation states in mammalian chromatin. Mol. Cell 12, 1577–1589.

65. Qin, X., Ahn, S., Speed, T.P., and Rubin, G.M. (2007). Global analyses of mRNA translational control during early Drosophila embryogenesis. Genome Biol. 8, 1–18.

66. Rämet, M., Pearson, A., Manfruelli, P., Li, X., Koziel, H., Göbel, V., Chung, E., Krieger, M., and Ezekowitz, R.A. (2001). Drosophila scavenger receptor CI is a pattern recognition receptor for bacteria. Immunity 15, 1027–1038.

67. Rämet, M., Manfruelli, P., Pearson, A., Mathey-Prevot, B., and Ezekowitz, R.A.B. (2002). Functional genomic analysis of phagocytosis and identification of a Drosophila receptor for E. coli. Nature 416, 644– 648.

68. Rea, S., Eisenhaber, F., O’Carroll, D., Strahl, B.D., Sun, Z.W., Schmid, M., Opravil, S., Mechtler, K., Ponting, C.P., Allis, C.D., et al. (2000). Regulation of chromatin structure by site-specific histone H3 methyltransferases. Nature 406, 593–599.

69. Redon, C., Pilch, D., Rogakou, E., Sedelnikova, O., Newrock, K., and Bonner, W. (2002). Histone H2A variants H2AX and H2AZ. Curr. Opin. Genet. Dev. 12, 162–169.

70. Ritzel, R.M., Patel, A.R., Grenier, J.M., Crapser, J., Verma, R., Jellison, E.R., and McCullough, L.D. (2015). Functional differences between microglia and monocytes after ischemic stroke. J. Neuroinflammation 12, 1–12.

71. Roddie, H.G., Armitage, E.L., Coates, J.A., Johnston, S.A., and Evans, I.R. (2019). Simu-dependent clearance of dying cells regulates macrophage function and inflammation resolution. PLoS Biol. 17, e2006741.

72. Ryglewski, S., Duch, C., and Altenhein, B. (2017). Tyramine Actions on Drosophila Flight Behavior Are Affected by a Glial Dehydrogenase/Reductase. Front. Syst. Neurosci. 11, 68.

73. Santos-Rosa, H., Schneider, R., Bannister, A.J., Sherriff, J., Bernstein, B.E., Emre, N.C.T., Schreiber, S.L., Mellor, J., and Kouzarides, T. (2002). Active genes are tri-methylated at K4 of histone H3. Nature 419, 407–411.

74. Schindelin, J., Arganda-Carreras, I., Frise, E., Kaynig, V., Longair, M., Pietzsch, T., Preibisch, S., Rueden, C., Saalfeld, S., Schmid, B., et al. (2012). Fiji: an open-source platform for biological-image analysis. Nat. Methods 9, 676–682.

75. Schmid, A., Chiba, A., and Doe, C.Q. (1999). Clonal analysis of Drosophila embryonic neuroblasts: neural cell types, axon projections and muscle targets. Development 126, 4653–4689.

76. Shepherd, D. (2000). Glial dependent survival of neurons in Drosophila. Bioessays 22, 407–409.

77. Sonnenfeld, M.J., and Jacobs, J.R. (1995a). Macrophages and glia participate in the removal of apoptotic neurons from the Drosophila embryonic nervous system. J. Comp. Neurol. 359, 644–652.

78. Sonnenfeld, M.J., and Jacobs, J.R. (1995b). Macrophages and glia participate in the removal of apoptotic neurons from the Drosophila embryonic nervous system. J. Comp. Neurol. 359, 644–652.

79. Stork, T., Engelen, D., Krudewig, A., Silies, M., Bainton, R.J., and Klambt, C. (2008). Organization and function of the blood-brain barrier in Drosophila. J. Neurosci. 28, 587–597.

80. Syed, A., and Tainer, J.A. (2018). The MRE11-RAD50-NBS1 Complex Conducts the Orchestration of Damage Signaling and Outcomes to Stress in DNA Replication and Repair. Annu. Rev. Biochem. 87, 263– 294.

81. Tepass, U., Fessler, L.I., Aziz, A., and Hartenstein, V. (1994). Embryonic origin of hemocytes and their relationship to cell death in Drosophila. Development 120, 1829–1837.

82. Thomas, P.D., Campbell, M.J., Kejariwal, A., Mi, H., Karlak, B., Daverman, R., Diemer, K., Muruganujan, A., and Narechania, A. (2003). PANTHER: A Library of Protein Families and Subfamilies Indexed by Function. Genome Res. 13, 2129–2141.

83. Trapnell, C., Pachter, L., and Salzberg, S.L. (2009). TopHat: discovering splice junctions with RNA-Seq. Bioinformatics 25, 1105–1111.

84. Volkenhoff, A., Weiler, A., Letzel, M., Stehling, M., Klämbt, C., and Schirmeier, S. (2015). Glial Glycolysis Is Essential for Neuronal Survival in Drosophila. Cell Metab. 22, 437–447.

85. Wang, Q., Goldstein, M., Alexander, P., Wakeman, T.P., Sun, T., Feng, J., Lou, Z., Kastan, M.B., and Wang, X.-F. (2014). Rad17 recruits the MRE11-RAD50-NBS1 complex to regulate the cellular response to DNA double-strand breaks. EMBO J. 33, 862–877.

86. Watts, R.J., Schuldiner, O., Perrino, J., Larsen, C., and Luo, L. (2004). Glia engulf degenerating axons during developmental axon pruning. Curr. Biol. 14, 678–684.

87. Wu, Y., Dissing-Olesen, L., MacVicar, B.A., and Stevens, B. (2015). Microglia: Dynamic Mediators of Synapse Development and Plasticity. Trends Immunol. 36, 605–613.

88. Zuber, R., Wang, Y., Gehring, N., Bartoszewski, S., and Moussian, B. (2020). Tweedle proteins form extracellular two-dimensional structures defining body and cell shape in Drosophila melanogaster. Open Biol. 10, 200214.

